# Mutant IDH silences GSX2 to reprogram neural progenitor cell fate and promote gliomagenesis

**DOI:** 10.1101/2025.08.12.667486

**Authors:** Yi Xiao, Diana D. Shi, Lei Guo, Ethan Neumann, Michael M. Levitt, Pranita Kaphle, Tracey Shipman, Haocheng Li, Feng Cai, Denise M. O. Ramirez, Lauren G. Zacharias, Zhenkang Chen, Mathew Lin, Vinesh T. Puliyappadamba, Tao Chen, Milan R. Savani, Salvador Peña, Janaka Wansapura, Thomas P. Mathews, Prashant Mishra, Yoon Jung Kim, Prithvi Raj, Timothy E. Richardson, Jian Xu, Stephen C. Mack, Gilbert J. Rahme, Bradley E. Bernstein, Ralph J. DeBerardinis, Itay Tirosh, Mario L. Suvà, Lin Xu, Kalil G. Abdullah, Samuel K. McBrayer

## Abstract

Isocitrate dehydrogenase (*IDH*) mutations arise early in gliomas and are associated with a defined neurodevelopmental cancer cell hierarchy. However, how mutant IDH contributes to this hierarchy and whether this interaction promotes gliomagenesis remain unclear. We captured the dynamics of IDH-mutant glioma initiation in genetically engineered mice through time-resolved, single-cell genomics. Mutant IDH activates and induces lineage switching of neural progenitor cells (NPCs). These actions expand oligodendrocyte precursor cells, the predominant cell-of-origin for these tumors, at the expense of interneurons. Lineage switching is mediated by promoter hypermethylation and silencing of *Gsx2*, a homeobox gene required for neurogenesis. Critically, *Gsx2* ablation recapitulates NPC fate reprogramming by mutant IDH. We provide a new model of neural cell fate control by *IDH* oncogenes and insights into the developmental origins of glioma.

## Main Text

Cancer-associated IDH mutations are recurrent in several human malignancies, including glioma, acute myeloid leukemia (AML), intrahepatic cholangiocarcinoma, and chondrosarcoma (*1*). IDH mutant enzymes synthesize the oncometabolite (*R*)-2-hydroxyglutarate [(*R*)-2HG] (*2*), which inhibits α-ketoglutarate-dependent TET and KDM dioxygenases that promote DNA and histone demethylation, respectively. Therefore, IDH-mutant cancer cells display chromatin hypermethylation phenotypes that promote oncogenesis (*3–10*).

In leukemias and cholangiocarcinomas, epigenetic reprogramming by (*R*)-2HG directly impacts cellular differentiation. Mutant IDH expression blocks hematopoietic cell differentiation (*3*) and enriches early hematopoietic progenitors (*11*) to promote leukemogenesis. Conversely, mutant IDH inhibition promotes lineage progression and terminal myeloid differentiation of hematopoietic cells in both preclinical leukemia models (*12*, *13*) and AML patients (*14*, *15*). In the liver, mutant IDH arrests hepatocyte differentiation and expands progenitor cells by silencing the lineage-specific transcription factor HNF-4α (*10*).

The impact of IDH mutations on neural cell differentiation in gliomas is less clear. *IDH* mutations occur early in glioma development (*16*) and define two histological subtypes of this disease: oligodendrogliomas and astrocytomas (*17*). In addition to *IDH* mutations, oligodendrogliomas are characterized by 1p/19q codeletion and *TERT* promoter mutations, whereas astrocytomas harbor *TP53* and *ATRX* mutations. Although astrocytomas and oligodendrogliomas have distinct genetic alterations, single cell genomics analyses of these tumors have established that they display comparable cellular hierarchies (*18–20*). Considered together, these findings imply that IDH oncogenes may play a conserved role in establishing this cellular hierarchy during glioma initiation. IDH-mutant gliomas comprise two broad classes of cells: 1) undifferentiated tumor cells, some of which are similar to NPCs (tri-potent cells capable of producing astrocytes, oligodendrocytes, and neurons), and 2) tumor cells displaying differentiation along astrocytic or oligodendroglial axes and reduced proliferative capacity. Although prior work has shown that mutant IDH activity controls astrocytic differentiation (*21–23*), we have limited understanding of the broader impact of IDH oncogenes on the cellular architecture of IDH-mutant gliomas.

Against this backdrop, two fundamental yet unresolved questions persist. Does mutant IDH control the fate of undifferentiated or glioma-initiating cell populations within this hierarchy? If so, what is the underlying molecular mechanism? Technical limitations have impeded efforts to answer these questions and unveil the origins of IDH-mutant glioma. Using patient samples to study this disease during premalignancy and tumor initiation is challenging because surgical specimens are acquired only after these phases of glioma evolution have passed. Additionally, expressing mutant IDH in certain genetic contexts paradoxically represses gliomagenesis in mice (*24*, *25*), which has impeded studies of the glioma-initiating activity of IDH oncogenes in preclinical models.

Here, we developed a genetic mouse model of mutant IDH-driven gliomagenesis and IDH-wildtype companion models to enable direct testing of causal genotype-phenotype relationships involving the *Idh1-R132H* oncogene. We leveraged these models to survey IDH-mutant glioma initiation by performing time-resolved, single-cell multiome (dual RNA and ATAC sequencing) analysis of engineered neural cells. We find that mutant IDH causes recurrent DNA hypermethylation and silencing of *Gsx2*, a gene required for neuroblast specification. By activating NPCs and blocking their neuronal differentiation, mutant IDH expands a population of oligodendrocyte precursor cells (OPCs) that give rise to IDH-mutant gliomas. Our data inform a conceptual framework that reveals how *IDH* oncogenes reshape neural cell differentiation trajectories during gliomagenesis.

## Results

### Establishing a highly penetrant genetic mouse model of IDH-mutant glioma

To investigate how mutant IDH initiates gliomagenesis, we sought to develop a genetic mouse model in which the *Idh1-R132H* oncogene promotes spontaneous glioma formation with nearly complete penetrance. We used recombinant AAV- and CRISPR/Cas9-based in vivo engineering to induce *Idh1* (*26*) and *Pik3ca* (*27*) gain-of-function mutations and *Trp53* and *Atrx* loss-of-function mutations in primitive, nestin^+^ cells of neonatal mice (**Fig. 1A and 1B**). *IDH1*, *TP53*, *ATRX* mutations commonly co-occur in human astrocytomas (*17*), while alterations in *PIK3R1* and *PIK3CA* genes that encode PI3K subunits are observed in a subset of these tumors (*28*, *29*). Somatic *IDH1* mutations display clonal expansion in some nondiseased human brain tissues (*30*). In tissue from an individual with an *IDH1-R132H*-mutant clone, this mutation was predominantly detected in a glia-enriched neural cell population but also observed at lower frequency in neuronal cells. This finding suggests that IDH mutations arise in tri-potent NPCs. Therefore, we elected to target our genetic engineering strategy toward nestin^+^ NPCs that give rise to both neuronal and glial lineages.

**Fig. 1.**
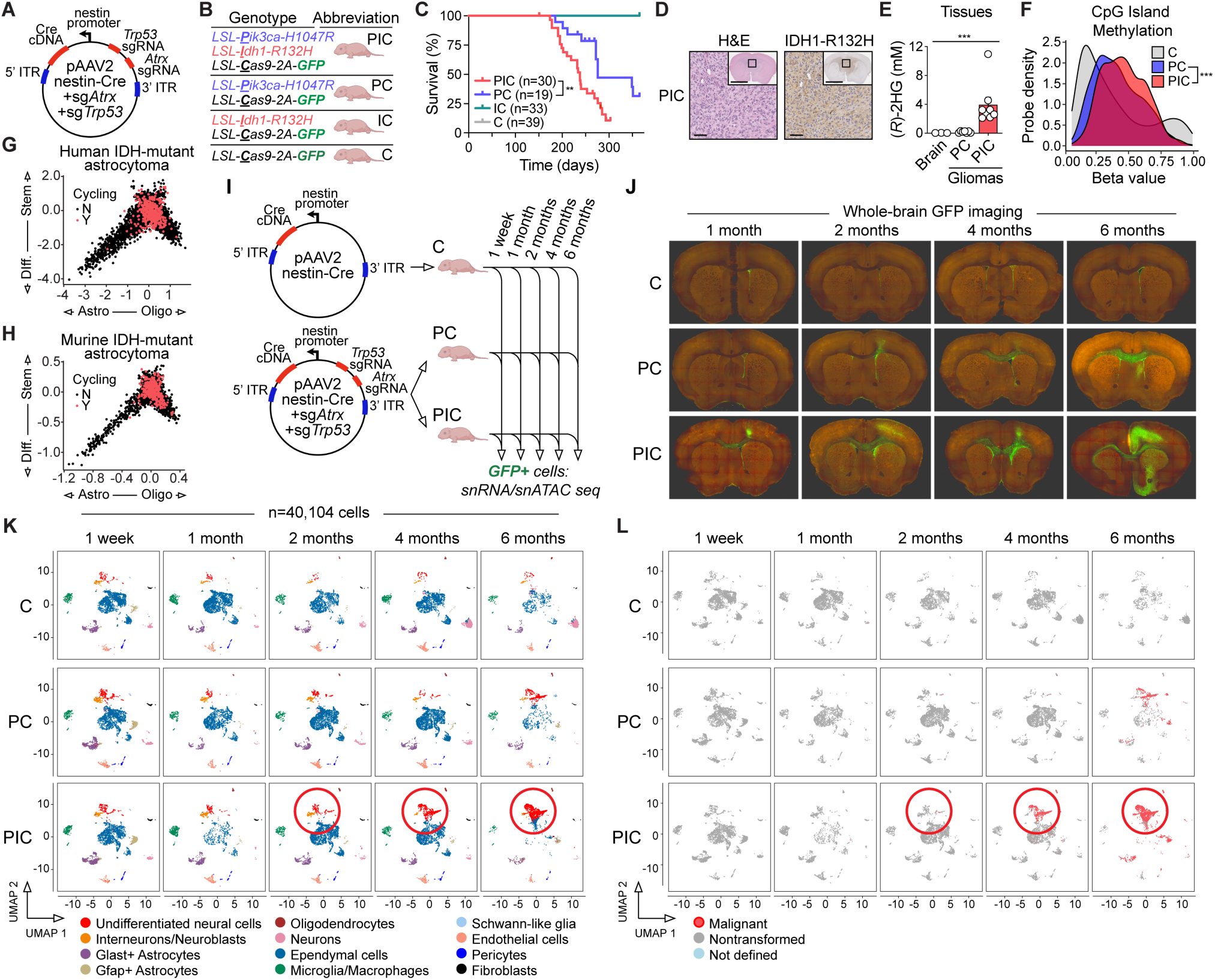
Time-resolved single-cell analysis of mutant IDH1-driven glioma initiation. (**A**) Diagram of AAV vector injected intracerebroventricularly into mouse strains in B. (**B**) Conditional compound transgenic mouse strains and abbreviations. LSL = loxP-stop-loxP. (**C**) Glioma-specific survival of mice of indicated genotypes injected intraventricularly with AAV vector in A. (**D**) H&E and IDH1-R132H IHC analyses of a representative astrocytoma in a PIC mouse. Scale bars: insets = 5 mm, main figures = 100 µm. Black boxes in insets indicate region in main figure. (**E**) (*R*)-2HG concentrations in non-malignant mouse brain tissue and in gliomas from PIC and PC mice. (**F**) CpG island methylation in non-malignant mouse brain tissue from C mice and in gliomas from PIC and PC mice. (**G**) Cell state analyses of human (Venteicher et al., Science, 2017) and (**H**) mouse (PIC mice) IDH-mutant astrocytomas. (**I**) Schema of experimental strategy. Note C mice express Cas9-2A-GFP but harbor no mutations and serve as a control for neurodevelopmental processes. (**J**) Whole-brain GFP imaging reveals spatial and temporal dynamics of GFP^+^ engineered neural cells in each cohort. (**K**) UMAP plots of cell state changes across genetic cohorts and time points. Plots were generated from integrated snRNAseq and snATACseq datasets collected from ≥3 mice at each time point in each cohort. Mice without detectable gliomas on MRI are shown at the 6-month time point in the PC cohort. (**L**) Single-nucleus copy number analysis using snRNAseq data. Malignant cells are positively identified via aneuploidy. Nontransformed cells are diploid. In (C), *P* value was calculated by log-rank test, ** *P* < 0.01. In (E), data are means ± SEM, *P* value was calculated by a one-way ANOVA nonparametric test, *** *P* < 0.001. In (F), *P* value was calculated by Kolmogorov-Smirnov (K-S) test, *** *P* < 0.001. In (K) and (L), red circles highlight mutant IDH-driven expansion and transformation of undifferentiated neural cell population.

To mutate these four genes, we first bred mice with inducible *Pik3ca* and *Idh1* oncogenes and an inducible *Cas9-2A-GFP* transgene (*LSL-<u>P</u>ik3ca-H1047R^+/-^*; *LSL-Idh1-R132H^+/wt^*; *LSL-<u>C</u>as9-2A-GFP^+/-^*, hereafter PIC mice). Two days after birth, transgenic pups were injected intracerebroventricularly with AAV particles expressing both a nestin enhancer-driven Cre cDNA (to activate oncogene and transgene expression) and sgRNAs targeting *Trp53* and *Atrx* genes. As controls, we also injected AAV particles into mice lacking one or both *Idh1* and *Pik3ca* oncogenes. These strains are referred to as PC (*LSL-<u>P</u>ik3ca-H1047R^+/-^*; *LSL-<u>C</u>as9-2A-GFP^+/-^*), IC (*LSL-<u>I</u>dh1-R132H^+/wt^*; *LSL-<u>C</u>as9-2A-GFP^+/-^*), and C (*LSL-<u>C</u>as9-2A-GFP^+/-^*) mice.

This work built on our past modeling efforts that produced IDH-mutant gliomas in some but not all mice (*28*, *31*). To achieve specific, uniform development of autochthonous IDH-mutant gliomas in mice, we incorporated a tissue-specific Cre driver and engineered mutations in younger mice. We also incorporated a Cre allele linked to GFP to track the fates of engineered cells and recover them for molecular analyses.

PIC mice developed gliomas with virtually full penetrance and displayed median overall survival of approximately 7.5 months (**Fig. 1C**). In comparison, PC mice (lacking mutant IDH1 but harboring cooperating mutations) displayed prolonged survival and reduced tumor incidence. IC and C mice lacking mutant PIK3CA did not develop gliomas. Therefore, the *Idh1-R132H* oncogene drives gliomagenesis in the context of cooperating *Pik3ca*, *Atrx*, and *Trp53* mutations in PIC mice. Gliomas that formed in PIC mice were reviewed by a board-certified neuropathologist (T.E.R.) and characterized as grade 3 astrocytomas (**Fig. 1D**). These tumors universally expressed the IDH1-R132H oncoprotein and accumulated pathophysiologically relevant (*R*)-2HG levels of ∼4 mM (*2*) (**Fig. 1D and 1E**). Glioma neurosphere lines established from tumors in PC and PIC mice displayed *Pik3ca* oncogene activation and mutational inactivation of *Trp53* and *Atrx* genes, while only the latter expressed mutant IDH1 (**Fig. S1A-S1R**). CRISPR indels in *Trp53* and *Atrx* genes were homogenous within each line, supporting a clonal origin of tumors in our model. Gliomas in PIC mice exhibited DNA hypermethylation (**Fig. 1F** and **Data S1**), recapitulating the CpG island methylator phenotype (CIMP) of human IDH-mutant gliomas (*4*). Further, the cellular hierarchy in PIC gliomas was reminiscent of that in human IDH-mutant astrocytomas (**Fig. 1G and 1H**) (*18*, *19*). Taken together, these findings indicate that the PIC mouse model of mutant IDH-driven astrocytoma reflects key histological, metabolic, epigenomic, and cellular features of the human disease.

### Time-resolved single-cell genomics analysis of glioma initiation

We sought to use this model to monitor the cellular and molecular dynamics of gliomagenesis and define effects that are causally linked to mutant IDH activity. To do so, we compared isogenic, engineered PIC and PC mice that differ only in *Idh1* mutational status (**Fig. 1I**). We also removed *Trp53* and *Atrx* sgRNAs from the AAV vector used for neural cell engineering and injected it into C mice. In this experiment, the only genetic alteration introduced in C mice entailed activation of the *Cas9-2A-GFP* transgene. Therefore, C mice served as controls to distinguish neurodevelopmental processes from those linked to gliomagenesis. We performed timed sacrifice of C, PC and PIC mice at defined intervals and harvested brain tissues for whole-brain GFP imaging, histology, immunohistochemistry, and single-nucleus dual RNA and ATAC sequencing analyses.

PIC mice displayed progressively increasing outgrowth and migration of engineered, GFP^+^ neural cells relative to PC and C cohorts, culminating in overt tumor development after 4 months (**Fig. 1J and S2A-S2M**). These differences cannot be explained by variability in AAV transduction efficiency between the cohorts because we observed GFP^+^ cells among the SOX2^+^ periventricular neural progenitors in all AAV-injected mice (**Fig. S2A-S2L**).

Next, we dissociated brain tissues from C, PC, and PIC mice sacrificed one week or one, two, four, or six months after AAV injection and enriched engineered, GFP^+^ neural cells by flow cytometry for single-cell genomics. In total, we profiled 40,104 cells by single-nucleus RNA and ATAC sequencing across all cohorts and time points (**Fig. 1K**). Our analysis captured expected neural cell populations, which were identified by canonical gene expression patterns (**Fig. S3**).

Notably, our dataset included large populations of ependymal cells owing to our intracerebroventricular AAV injection strategy. We also recovered cells that are not derived from nestin^+^ neural precursors (including myeloid cells, pericytes, and endothelial cells), reflecting contaminating, GFP^-^ cells retained during sample processing and/or low frequency Cre activation in these cell populations.

Comparing cell populations over time revealed a specific increase in undifferentiated neural cells in PIC mice relative to PC and C. This population expressed markers of proliferative or primitive neural cell states, including *Top2a*, *Cenpf*, *Pdgfra*, *Olig2*, and *Sox2* (**Fig. S3B**). This trend began at one-month post-AAV injection and peaked at six months, at which time this population became the largest of those detected. In C mice, this population steadily declined, reflecting the differentiation of stem and progenitor cells in the CNS after birth. A similar trend was observed in PC mice over the first four months, although this population expanded at the end of this timecourse. Because these populations reflected engineered cells spanning many neural cell types, we could directly and concurrently compare their permissiveness to transformation by the *Idh1-R132H* oncogene and cooperating mutations. We used single-nucleus RNA sequencing data to infer copy number alterations in individual cells and categorized them as either aneuploid and malignant or diploid and nontransformed (**Fig. 1L**), consistent with prior studies (*18*, *32*). This strategy was feasible because engineered murine gliomas, like analogous human gliomas (*33*), commonly displayed tumor-specific copy number alterations. We observed virtually no aneuploid cells in C mice lacking oncogenic mutations across all time points. In PIC mice, the malignant clone emerged within the undifferentiated neural cell population between two and four months after genetic engineering.

### Mutant IDH reprograms NPC fate, leading to expansion and transformation of OPCs

We next examined changes in undifferentiated neural cells and the closely related population of neuroblasts and interneurons. Focused UMAP dimensionality reduction analysis revealed that undifferentiated neural cells comprised two populations, neural progenitor cells (NPCs) and oligodendrocyte precursor cells (OPCs) (**Fig. S4A-I**). NPCs displayed high expression of cell cycle-related genes compared with other cell populations, in keeping with a cellular hierarchy in which NPC division and subsequent differentiation generates either neuronal or glial progeny.

Neural cells harvested from C mice engineered with nestin-Cre only (i.e. functionally wild-type) primarily displayed NPC or interneuron/neuroblast, but not OPC, identity at 1 month of age and older (**Fig. 2A and 2B**). These findings are consistent with canonical cellular differentiation patterns in the postnatal subventricular zone (SVZ): NPCs principally generate neuroblasts and interneurons that migrate away from the SVZ in defined patterns throughout the brain (*34*).

**Fig. 2.**
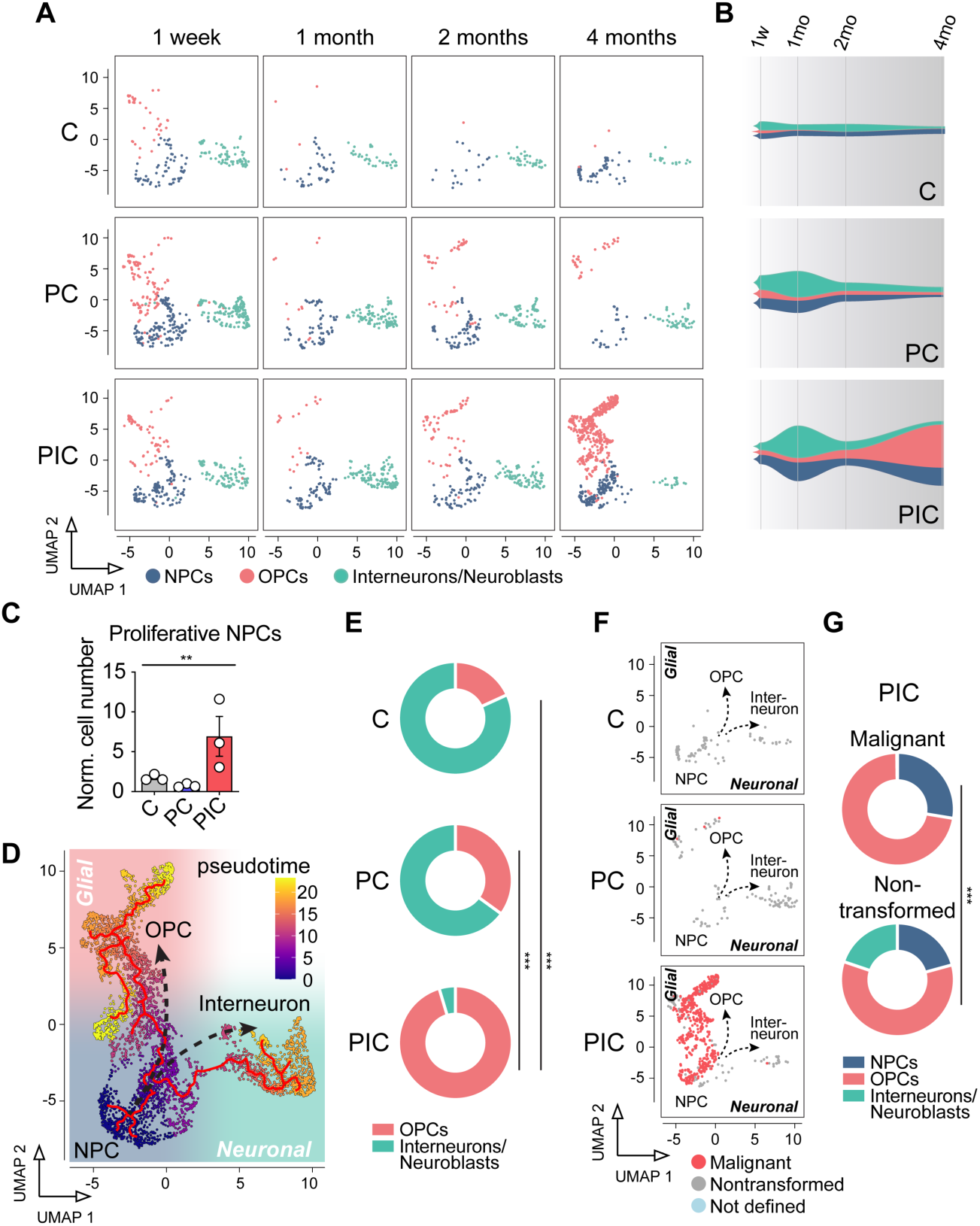
Mutant IDH activates and induces lineage switching of NPCs during glioma initiation. (**A**) UMAP plots of cell populations within “undifferentiated neural cell” and “interneuron/neuroblast” cell states from data in Figure 1K. (**B**) Fish plots showing cell population dynamics modeled from single-nucleus data in (A). (**C**) Quantification of proliferative NPCs in C, PC, and PIC mice at 4 months. Cell numbers were normalized (Norm.) by calculating their incidence per 100 cells sequenced in GFP^+^ populations. (**D**) Pseudotime analysis of data in (A) showing neuronal and glial differentiation trajectories of NPCs overlaid on combined UMAP plot. (**E**) Relative abundance of OPCs and interneurons/neuroblasts at 4 months. (**F**) Single-nucleus copy number analysis of NPCs, OPCs, and interneurons/neuroblasts from PIC mice at 4 months using snRNAseq data. Malignant cells are positively identified via aneuploidy. Nontransformed cells are diploid. (**G**) Relative quantification of cell states within malignant and nontransformed cell populations determined in (F). In (C), data are means ± SEM, *P* value was calculated by a one-way ANOVA nonparametric test, ** *P* < 0.01. In (E) and (G), *P* values were calculated by Chi-square tests, *** *P* < 0.001.

Trends were similar in PC mice, except for a transient increase in NPC and interneuron/neuroblast numbers after induction of *Pik3ca*, *Trp53*, and *Atrx* mutations. In contrast, cell population dynamics were substantially altered by mutant IDH activity. Comparing PIC with PC and C mice showed that mutant IDH caused expansion of OPCs and NPCs and contraction of neuroblasts and interneurons in the months leading up to malignant transformation. Using gene expression data to infer proliferative status of individual cells (**Fig. S4J**), we determined that these effects were associated with an increased number of cycling NPCs in PIC mice (**Fig. 2C**). We inferred cellular trajectories via pseudotime analysis and found that NPCs differentiate along two mutually exclusive axes. NPCs either display glial lineage priming to generate OPCs or adopt a neuronal fate and sequentially produce neuroblasts and interneurons (**Fig. 2D**). Notably, we did not observe a trajectory through which glial-primed progenitors are reprogrammed to a neuronal fate or vice versa. Mutant IDH alters NPC fate decision, promoting OPC differentiation at the expense of neuronal lineages (**Fig. 2E and S4K-M**). Taken together, these data demonstrate that mutant IDH activates NPCs and switches the fate of their progeny from a neuronal to a glial lineage. These effects increase relative (**Fig. 2E**) and absolute (**Fig. S4N-Q**) numbers of OPCs and NPCs.

NPC lineage switching caused by mutant IDH1 is functionally relevant, because neuroblast and interneuron cell states were incompatible with malignant transformation by the *Idh1-R132H* oncogene and cooperating mutations (**Fig. 2F, 2G, and S4R-T**). Rather, the malignant clone that emerged in PIC mice was primarily associated with the OPC cell state (**Fig. 2F and 2G**). IDH-mutant OPCs displayed low proliferative potential prior to transformation (**Fig. S4U**), indicating that mutant IDH-dependent expansion of OPCs in premalignancy is largely driven by NPC fate reprogramming rather than aberrant cycling of OPCs.

### Mutant IDH represses Gsx2 and an interneuron developmental gene expression program

To discover the molecular mechanism by which mutant IDH causes NPC lineage switching, we conducted a cross-species transcriptomic analysis. We first performed bulk RNA-sequencing analysis of engineered neural cells isolated from PIC and PC mice (**Data S2**). We found that mutant IDH activity suppressed genes related to interferon signaling (**Fig. S5A-E and Data S3**), aligning with results of our (*9*, *35*) and others’ prior work (*36*) and validating this dataset. Next, we identified transcripts that are differentially expressed between human IDH1/2-mutant and IDH1/2-wildtype gliomas and between cells from PIC and PC mice (**Fig. 3A**). We focused our analysis on genes with known roles in gliogenesis or neurogenesis, or those that encode transcription factors (**Data S4**). Using this curated list, we found that multiple genes involved in interneuron differentiation were downregulated in both human and murine IDH-mutant samples, including *GSX2*, *DLX1*, *DLX2*, *SP8*, and *SP9* (**Fig. 3B**). GSX2 acts as a master regulator of neuronal versus glial cell fate decisions in ganglionic eminence- and SVZ-resident NPCs during prenatal and postnatal brain development, respectively (*40*, *44*). When activated, this lineage-specific transcription factor promotes neuroblast lineage commitment, inhibits expression of genes involved in glial differentiation (*37*, *38*), and primes the expression of transcription factors required for interneuron development, including DLX1, DLX2, SP8, and SP9 (*39*). Consistent with this model, we found that *Dlx1*, *Dlx2*, *Sp8*, and *Sp9* are nearly exclusively expressed in NPC and neuroblast/interneuron populations while *Gsx2* is expressed largely by NPCs that are poised for glial or neuronal differentiation (**Fig. S4A and S6A-S6E**). Although *Gsx1*, a paralog of *Gsx2*, has also been implicated in neural cell fate regulation (*40*, *41*), it was nearly undetectable in all cell populations (**Fig. S6F**). Therefore, *Gsx1* was not studied further.

**Fig. 3.**
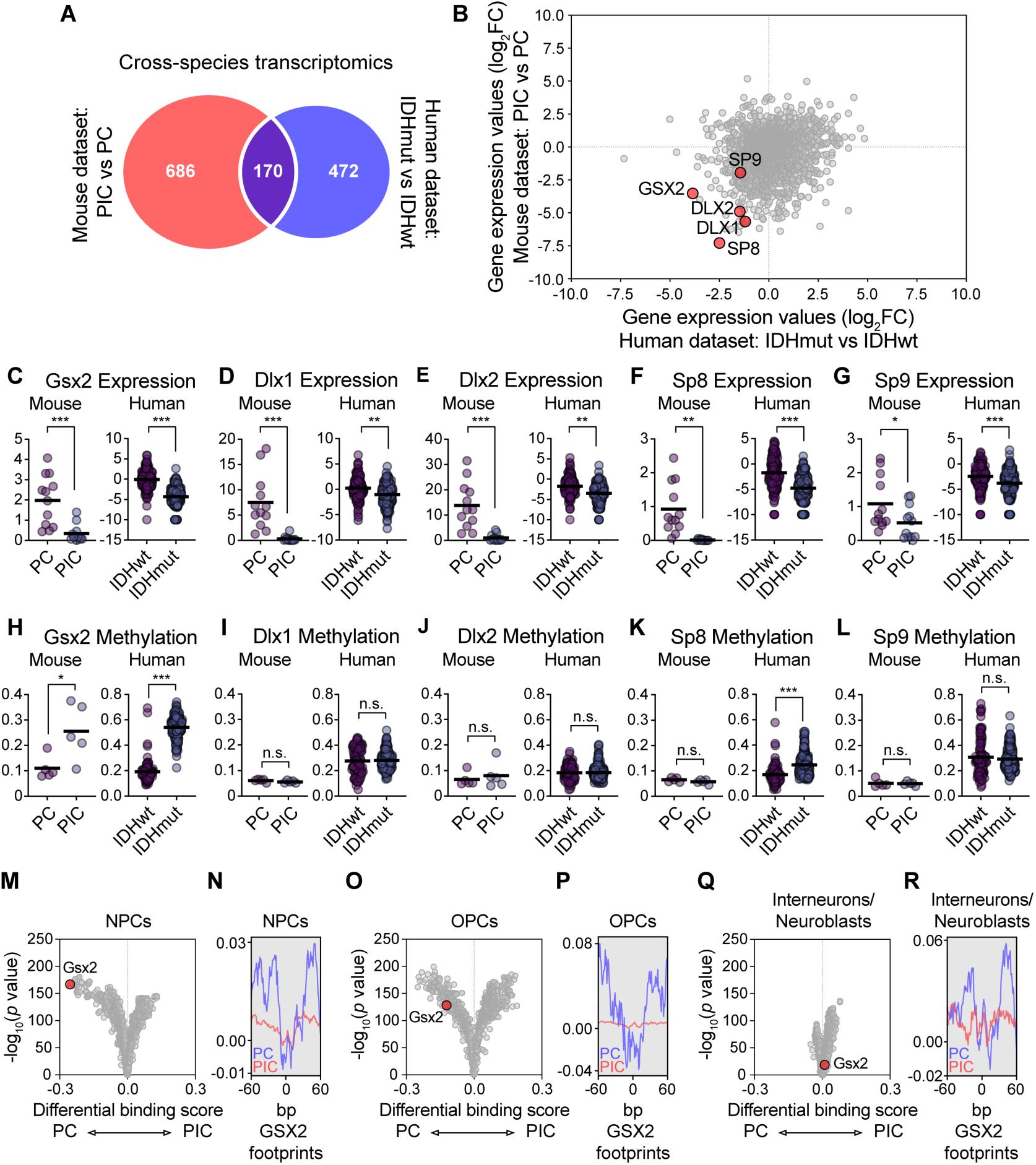
Repression of neuroblast and interneuron gene expression programs by IDH mutations. (**A**) Venn diagram of differentially expressed genes with known roles in gliogenesis or neurogenesis or those that encode transcription factors between murine (PIC versus PC mice at 6 months) and human (TCGA GBMLGG dataset) IDH-mutant (IDHmut) and IDH-wildtype (IDHwt) brain tissues. (**B**) Scatter plot showing differentially expressed interneuron lineage genes in IDHmut versus IDHwt murine and human brain tissues. (**C**)-(**L**) Interneuron lineage gene (C)-(G) expression and (H)-(L) DNA methylation in IDHmut versus IDHwt murine and human brain samples. For expression data, y-axes values represent log_2_(FPKM + 0.001) for human and FPKM for mouse. For DNA methylation data, y-axes values are beta values. (**M**)-(**R**) Comparison of genome-wide DNA binding scores for transcription factors in NPCs (M)-(N), OPCs (O)-(P), and interneurons/neuroblasts (Q)-(R) from PIC and PC mice at 4 months. In (M), (O), and (Q), volcano plots are shown, with the TOBIAS differential binding score on the x-axis and -log_10_(p value) on the y-axis. In (N), (P), and (R), footprint plot overlays are shown for GSX2 in cells from PIC (red line) and PC (blue line) mice. In (C)-(L), bars are mean values; *P* values were calculated by unpaired t-tests, * *P* < 0.05, ** *P* < 0.01, *** *P* < 0.001, n.s. = not significant.

We hypothesized that mutant IDH may silence one or more of these genes via DNA hypermethylation, given that (*R*)-2HG inhibits TET 5’ methylcytosine hydroxylases that promote genomic DNA cytosine demethylation. We directly compared expression and DNA methylation (**Data S1**) of *GSX2*, *DLX1*, *DLX2*, *SP8*, and *SP9* genes in murine and human IDH-mutant and IDH-wildtype samples. Although each gene was significantly downregulated in IDH-mutant samples (**Fig. 3C-3G**), only *GSX2* was hypermethylated in IDH-mutant samples from both species (**Fig. 3H-3L**). Neither downregulation of this gene set nor *GSX2* hypermethylation was observed in other IDH-mutant human cancers, including cholangiocarcinoma and acute myeloid leukemia (**Fig. S7**). Cancer-associated loss-of-function mutations affecting succinate dehydrogenase (SDH) or fumarate hydratase (FH) enzymes cause accumulation of succinate and fumarate. These oncometabolites, like (*R*)-2HG, inhibit TET enzymes and cause CIMP in tumors (*42*, *43*). Although SDH- and FH-mutant cancers displayed DNA hypermethylation of these interneuron-related genes in several instances, none were downregulated in these tumors relative to SDH- or FH-wildtype counterparts (**Fig. S8**). Therefore, *GSX2* hypermethylation and downregulation of interneuron developmental genes are specific features of IDH-mutant gliomas.

Our findings suggest that GSX2-dependent gene transactivation may be repressed in IDH-mutant neural cells. To test this idea, we performed a genome-wide transcription factor footprinting analysis using single-cell ATAC-sequencing data generated from PIC, PC, and C engineered neural cells (**Data S5**). Compared with genome binding profiles of PC and C cells, GSX2 occupancy was strongly attenuated in both NPC and OPC cells from PIC mice (**Fig. 3M-P and S9A-S9D**). Among all transcription factors, GSX2 exhibited one of the most pronounced reductions in activity in NPCs from PIC mice relative to PC mice (**Fig. 3M and 3N**). In contrast, GSX2 binding was not substantially altered in interneuron and neuroblast populations (**Fig. 3Q, 3R, S9E, and S9F**). These findings are consistent with GSX2 being activated in NPCs during neuroblast specification and subsequently silenced as neuronal differentiation progresses (*45*).

### Promoter CpG island hypermethylation is sufficient to explain Gsx2 silencing by mutant IDH

We explored the relationship between *GSX2* DNA methylation and expression in more detail. In both mouse and human genomes, *GSX2* is associated with large, promoter-spanning CpG islands (**Fig. 4A**). It is plausible that *GSX2* is particularly susceptible to (*R*)-2HG-evoked DNA hypermethylation due to its high CpG content. This idea is supported by gene set enrichment analysis showing that genes that have high CpG (HCP) content and are marked by H3K27 trimethylation in NPCs (including *Gsx2*) are downregulated in PIC versus PC cells (**Fig. 4B**).

**Fig. 4.**
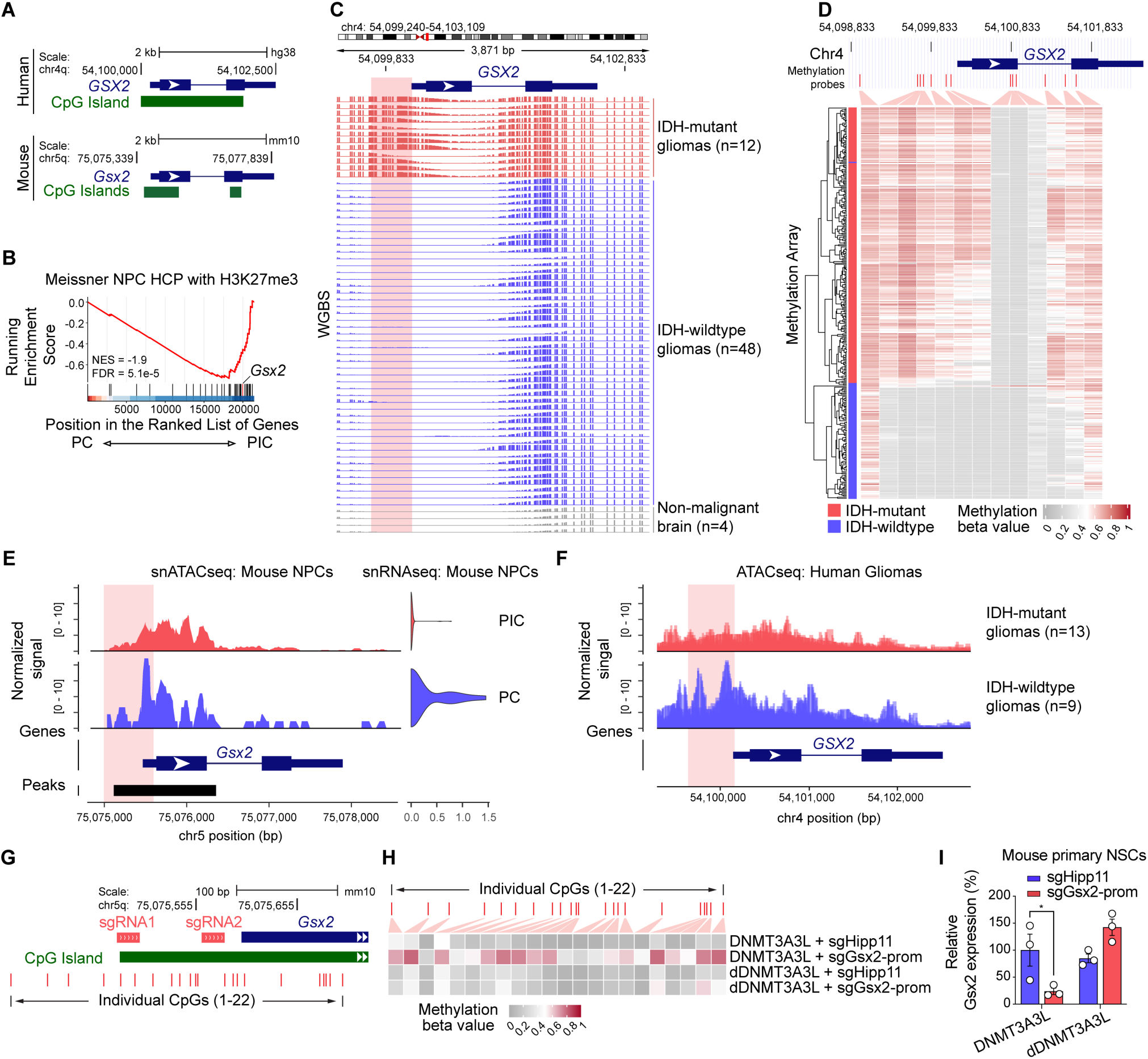
NPC lineage switching is associated with promoter hypermethylation and silencing of the homeobox gene *Gsx2*. (**A**) Schema of human *GSX2* and mouse *Gsx2* genes and associated CpG islands. (**B**) GSEA of engineered cells from PIC and PC mice. “Meissner NPC HCP with H3K27me3” gene set encompasses genes with high CpG (HCP) content and H3K27me3 marks in NPCs, including *Gsx2* (red line). (**C**) Whole genome bisulfite sequencing tracks of human glioma samples and non-malignant brain tissues show hypermethylation of the *GSX2* promoter region in IDHmut gliomas compared to IDHwt gliomas and non-malignant brain specimens. (**D**) Heatmap of methylation array data for all probes associated with *GSX2* from the TCGA GBMLGG cohort. (**E**) Left: coverage plot of aggregated single-nucleus ATAC-seq peaks at the *Gsx2* locus in NPCs from PIC and PC mice at 4-month time point. Right: *Gsx2* expression in NPCs in from PIC and PC mice at 4-month time point. (**F**) Coverage plot of ATAC-seq peaks from human gliomas at the *GSX2* locus. (**G**) Schema of two CRISPR sgRNAs targeting the mouse *Gsx2* promoter region and associated CpG islands. (**H**) Heatmap showing methylation levels of 22 CpGs in the mouse *Gsx2* promoter in primary mNSCs. Cells were transfected with the dCas9-DNMT3A3L epigenome editing construct and *Gsx2*-targeting sgRNAs or with control constructs (*Hipp11* safe harbor locus targeting sgRNAs or catalytically dead dCas9-dDNMT3A3L). (**I**) Relative quantification of *Gsx2* expression by real-time qRT-PCR in primary mNSCs from (H). In (I), data are means ± SEM; *P* value was calculated by a two-way ANOVA with Fisher’s least significant difference (LSD) test. * *P* < 0.05.

This finding was broadly applicable to HCP genes with different patterns of histone methylation (**Fig. S10A-C and S10E**), except for those with bivalent domains featuring both H3K4me3 and H3K27me3 marks (**Fig. S10D**). In contrast, genes with low CpG (LCP) content displayed a comparatively weak trend toward reduced expression in PIC versus PC cells that did not reach statistical significance (**Fig. S10F**).

Assessing DNA methylation in two human tissue datasets revealed pervasive, specific methylation of the CpG island associated with the *GSX2* promoter in IDH-mutant gliomas relative to IDH-wildtype gliomas and non-malignant brain (*46*) (**Fig. 4C and 4D**). The association between this epigenetic feature and *IDH1/2* genotype was remarkably strong, as unsupervised hierarchical clustering of samples according to *GSX2* methylation nearly perfectly separated IDH-mutant and IDH-wildtype gliomas (**Fig. 4D**). *GSX2* promoter hypermethylation correlated with decreased chromatin accessibility at this locus in both PIC NPCs and human IDH-mutant gliomas relative to their IDH-wildtype counterparts (**Fig. 4E and 4F**). These data establish that promoter hypermethylation of *GSX2* is a conserved consequence of mutant IDH activity in glioma.

To ask if promoter hypermethylation is sufficient to cause *Gsx2* silencing, we performed CRISPR-based epigenome editing in primary mouse neural stem cells (mNSCs). After validating relevant expression constructs in HEK293 cells (**Fig. S11**), we transfected mNSCs to express inactive Cas9 nuclease (dCas9) fused to either an active (DNMT3A/3L) or catalytically dead (dDNMT3A/3L) DNA methyltransferase (*47*). Simultaneously, we expressed a pair of sgRNAs targeting the *Gsx2* promoter (**Fig. 4G**) or the *Hipp11* safe harbor locus. mNSCs expressing dCas9-DNMT3A3L and *Gsx2* sgRNAs displayed both *Gsx2* hypermethylation (**Fig. 4H**) and downregulation (**Fig. 4I**) relative to negative controls. Therefore, mutant IDH-induced promoter hypermethylation is sufficient to explain *Gsx2* downregulation.

### Gsx2 deficiency is sufficient to recapitulate NPC lineage switching caused by mutant IDH

We next investigated how GSX2 activity during premalignancy affects the generation of OPCs, the predominant tumor-initiating cell population in IDH-mutant gliomas. To do so, we modified the AAV vector used for in vivo neural cell engineering (**Fig. 1A**), adding sgRNAs targeting either *Gsx2* or the *Hipp11* safe harbor locus (**Fig. 5A**). To study the consequences of *Gsx2* silencing independent of other effects of mutant IDH activity, we introduced these AAV particles into IDH-wildtype PC mice via intracerebroventricular injection. When PC mice reached two or three months of age, we sacrificed them, isolated GFP^+^ engineered neural cells from dissociated brain tissues, and performed single-cell RNA sequencing. At two months of age, mice injected with AAV particles expressing *Hipp11* or *Gsx2* sgRNAs did not display significant differences in the relative sizes of NPC, OPC, or interneuron/neuroblast populations (**Fig. 5B and 5C**) despite evidence of robust CRISPR indel formation in the *Gsx2* gene in the latter (**Fig. S12**). In both cohorts, interneurons and neuroblasts outnumbered OPCs. However, at three months, *Gsx2* deficiency caused an inversion in the relative sizes of these cell populations, with engineered OPCs outnumbering interneurons and neuroblasts. A similar effect was not observed in the control cohort at this time point. The effect size of lineage switching caused by *Gsx2* knockout (**Fig. 5D**) mirrored that caused by mutant IDH1 prior to malignant transformation (**Fig. 5E**).

**Fig. 5.**
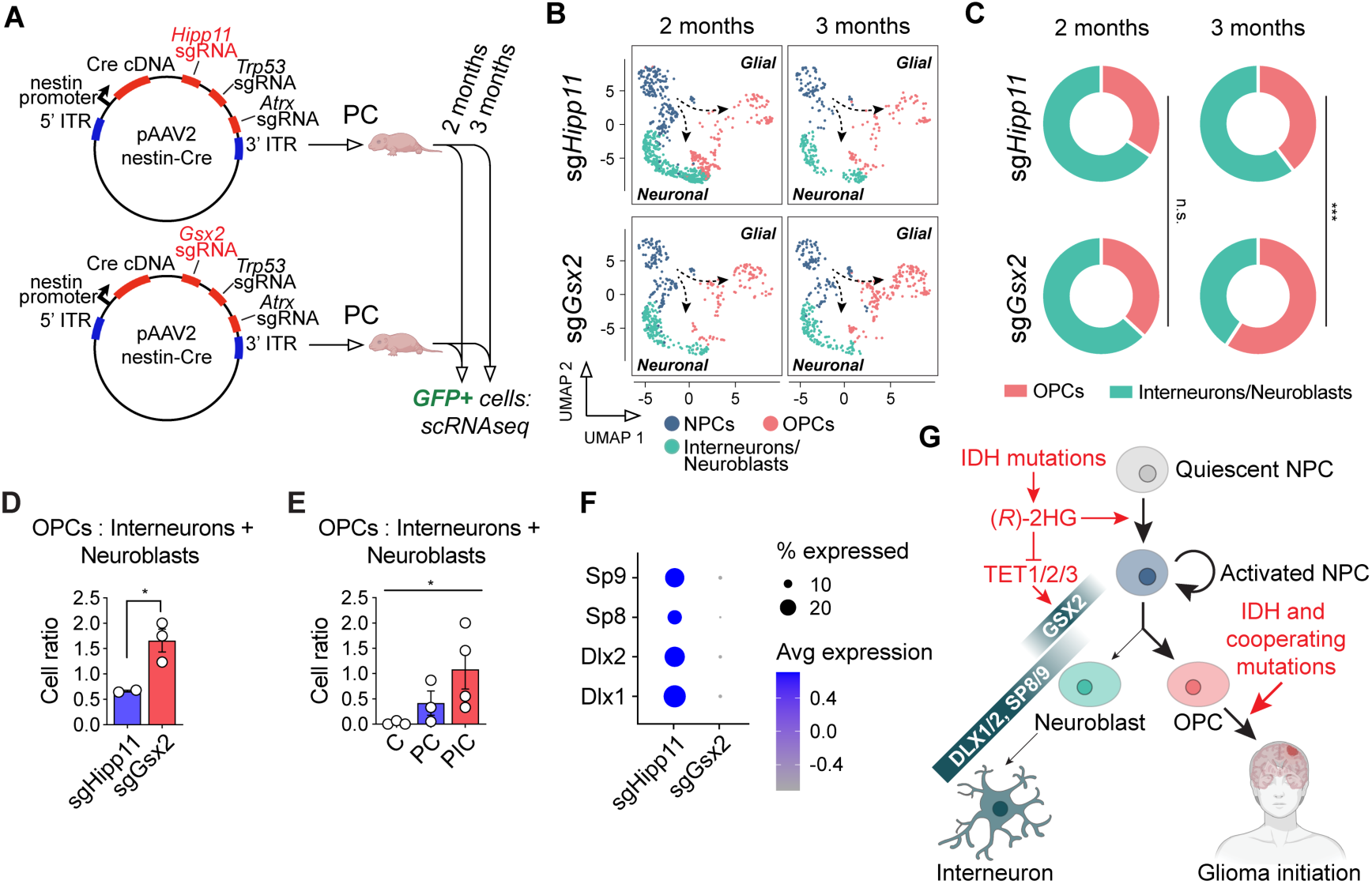
*Gsx2* knockout recapitulates NPC fate reprogramming by mutant IDH. (**A**) Schema of experimental strategy. PC mice were injected with modified AAV vector with an additional sgRNA targeting either the *Hipp11* safe harbor locus or *Gsx2*. (**B**) UMAP plots showing NPCs, OPCs, and interneurons/neuroblasts recovered from PC mice 2 or 3 months post-injection with AAVs encoding *Hipp11* or *Gsx2* sgRNAs. (**C**) Relative abundance of OPCs and interneurons/neuroblasts calculated from data in (B). (**D**) Ratio of OPCs to interneurons and neuroblasts calculated from data in (B). (**E**) Ratio of OPCs to interneurons and neuroblasts recovered from PIC, PC, or C mice at 2 months. (**F**) Expression of interneuron lineage genes in cells from PC mice engineered with *Hipp11* or *Gsx2* sgRNAs. (**G**) Model linking IDH mutations with NPC lineage switching via *GSX2* silencing. In (C), *P* values were calculated by Chi-square tests, *** *P* < 0.001, n.s. = not significant. In (D) and (E), data are means ± SEM; * *P* < 0.05. In (D), *P* value was calculated by an unpaired t-test. In (E), *P* value was calculated by a one-way ANOVA nonparametric test.

Interestingly, *Gsx2* knockout also inhibited expression of *Dlx1*, *Dlx2*, *Sp8*, and *Sp9*, recapitulating the impact of mutant IDH (**Fig. 3B-3G and 5F**). Therefore, loss of GSX2 expression is sufficient to explain mutant IDH-dependent repression of this interneuron gene expression program. Taken together, our findings inform a new model of mutant IDH action during glioma initiation (**Fig. 5G**). We propose that by activating NPCs and impairing GSX2-dependent neuroblast differentiation, IDH oncogenes enrich and transform OPCs in the presence of cooperating mutations.

## Discussion

These findings deepen our understanding of the intimate ties between IDH oncogenes and altered cell fate control in cancer. Seminal studies of IDH-mutant leukemia and cholangiocarcinoma established the ability of mutant IDH to arrest cellular differentiation (*3*, *10*, *12*, *14*, *15*). Recent work showing that mutant IDH inhibition induces astrocytic differentiation in human gliomas indicates that this property extends to brain tumors as well and highlights the therapeutic relevance of cellular hierarchies in glioma (*21*). Our results reveal that in addition to blocking differentiation, IDH mutations alter cell states in gliomas via another key mechanism: lineage switching. When activated, NPCs in the SVZ face a cell fate decision: differentiating along mutually exclusive neuronal or glial axes (*34*). We find that mutant IDH exerts two primary effects on the fate of NPCs in the postnatal brain. In agreement with prior work (*48*), mutant IDH activates and triggers outgrowth of NPCs. Mutant IDH also alters NPC fate decisions, driving glial differentiation at the expense of neuronal specification. Together, these two mechanisms are associated with OPC expansion during IDH-mutant glioma initiation.

NPC lineage switching and consequent enlargement of the OPC pool have important implications for gliomagenesis. The cell-of-origin for adult-type diffuse gliomas displays hallmarks of immature glial, rather than neuronal, cell states (*49–51*) and multiple lines of evidence support OPCs as the relevant cell-of-origin in IDH-mutant gliomas. The gene expression program that defines human IDH-mutant gliomas (*52*) closely recapitulates the transcriptional signature of OPCs (*18*, *19*). Moreover, recent studies analyzing rare IDH-mutant clones in large collections of human brain tissues implicated glia (*30*) and, more specifically, OPCs (*53*) as the source of tumor-initiating cells in gliomas that bear these mutations.

Although OPCs have been implicated as the cell-of-origin in this disease, one study of IDH-mutant clones in nondiseased brain tissue reported detection of the *IDH1-R132H* mutation in both glial-enriched and neuronal cell population, albeit at much higher frequency in the former (*30*). This suggests that IDH mutations first arise in neural precursors with the potential to generate both glia and neurons cells. Considering these findings together with our data, we propose a multi-step process of IDH-mutant glioma initiation. First, *IDH* mutations occur in tri-potent NPCs. Second, mutant IDH activity enhances proliferation and represses neuronal specification of NPCs. Third, IDH-mutant OPCs expand in number, culminating in emergence of the malignant clone principally from this transformation-permissive cell population. We modeled this process by engineering the *Idh1-R132H* oncogene and cooperating mutations into nestin^+^ primitive neural cells, including NPCs. Differentiation mechanisms introduced these mutations into distinct neural cell populations over time. Therefore, our approach yielded, in essence, an in vivo competition assay that allowed us to map cells comprising the malignant clone onto cell states at the time of transformation. Although both NPCs and OPCs can be transformed by these genetic alterations, our work provides direct evidence that OPCs represent the predominant cell-of-origin in IDH-mutant glioma. In contrast, we discovered that neuronal lineages are strictly incompatible with transformation by this set of oncogenes. This finding is specific to IDH-mutant glioma, because interneuron progenitors represent the cell-of-origin for histone H3.3G34-mutant diffuse hemispheric gliomas (*45*, *54*).

Mechanistically, we show that NPC lineage switching in murine IDH-mutant gliomas can be explained by epigenetic silencing of the homeobox gene *Gsx2*. We observed pervasive hypermethylation of a large CpG island spanning the *Gsx2* promoter in both murine and human IDH-mutant gliomas. This effect is likely attributable to (*R*)-2HG-dependent inhibition of the TET 5’methylcytosine hydroxylases that promote DNA demethylation (*3*, *55*), which underpins the CIMP in these tumors (*4*, *23*). In support of this model, we showed that experimentally inducing *Gsx2* promoter hypermethylation in IDH-wildtype NSCs was sufficient to repress *Gsx2*. *Gsx2* knockout depletes interneurons and causes OPCs to accumulate in the brains of prenatal and postnatal mice (*40*, *44*, *56*). Therefore, we hypothesized that GSX2 silencing may explain how mutant IDH reprograms NPCs to adopt a glial rather than neuronal fate. Remarkably, phenotypes associated with GSX2 suppression in our tumor models recapitulated those in normal neurodevelopment: both IDH-mutant PIC mice and *Gsx2*-knockout PC mice displayed OPC expansion and depletion of interneurons and neuroblasts. These findings may help explain a perplexing feature of the cellular architecture of human IDH-mutant gliomas (*18–20*). Although these tumors comprise cells that resemble tri-potent NPCs that should give rise to neuronal progeny, neuronal tumor cell states are largely absent from this architecture. Our data suggest that mutant IDH-dependent *Gsx2* silencing and NPC lineage switching contribute to this architecture by blocking neuroblast specification.

Despite intense investigation of IDH-mutant gliomas since their initial discovery (*57*), knowledge of the precise molecular mechanisms that link IDH mutations with oncogenesis is still accruing. Our findings provide new insights into this oncogenic program by revealing *GSX2* as a functionally relevant target of mutant IDH-dependent epigenetic reprogramming. Our work builds on studies demonstrating that DNA hypermethylation caused by (*R*)-2HG accumulation causes dysregulation of *PDGFRA*, *CDKN2A*, *CGAS*, and endogenous retroviruses (ERVs) (*6*, *8*, *9*), each of which are linked to tumor-promoting processes at play in IDH-mutant gliomas. *IDH* oncogenes also drive tumorigenesis through chromatin-independent mechanisms, including (*R*)-2HG-dependent activation of the EGLN1 prolyl hydroxylase and consequent degradation of the glial tumor suppressor HIF1α (*58*, *59*). As a full portrait of the oncogenic mechanisms evoked by IDH mutations in glioma emerges, evaluating potential cooperation between individual effector pathways is warranted.

Knowledge of the fundamental mechanisms of mutant IDH-dependent transformation hold increasing clinical relevance. Recently, the dual mutant IDH1/2 inhibitor vorasidenib was approved by the U.S. Food and Drug Administration to treat IDH-mutant low grade gliomas on the basis of positive phase 3 clinical trial results (*60*). Fundamental studies of the cellular processes and molecular events through which IDH mutations cause glioma initiation are vital, as they may inform rational strategies to monitor or predict responsiveness to mutant IDH inhibitors and develop combination treatment protocols involving this new class of glioma therapeutics.

## Acknowledgments

We thank members of the McBrayer, Abdullah, and DeBerardinis laboratories and Dr. Sean Morrison for helpful feedback; Dr. Matthew Garrett for sharing the human glioma ATAC-seq data; the Children’s Research Institute’s Flow Core Facility at UT Southwestern for assistance with cell sorting; and the UT Southwestern Animal Resource Center for assistance with in vivo studies in mice.

## Author contributions

Conceptualization: YX, SKM

Methodology: YX, DDS, LG, FC, DMOR, LGZ, ZC, TC, JW, TPM, PM, YJK, PR, JX, SCM, GJR, BEB, RJD, IT, MLS, LX, SKM

Investigation: YX, DDS, LG, EN, MML, PK, TS, HL, FC, DMOR, ML, VTP, TC, SP, JW, TER, SKM

Visualization: YX, LG, DMOR, LX, SKM

Funding acquisition: YX, KGA, SKM

Project administration: SKM

Supervision: KGA, SKM

Writing – original draft: YX, DDS, SKM

Writing – review & editing: YX, DDS, MRS, SCM, GJR, BEB, KGA, SKM

## Data and materials availability

All data are available in the manuscript or the supplementary materials. Further information and requests for resources and reagents should be directed to and will be fulfilled by the corresponding author, Samuel K. McBrayer. Omics data will be deposited Gene Expression Omnibus (GEO) website.

## Supplementary Materials

Materials and Methods

Figs. S1 to S12

References (*1–22*)

Data S1 to S5

## References

1. J. A. Losman, W. G. Kaelin, What a difference a hydroxyl makes: mutant IDH, (R)-2-hydroxyglutarate, and cancer. Genes Dev 27, 836–52 (2013).

2. L. Dang, D. W. White, S. Gross, B. D. Bennett, M. A. Bittinger, E. M. Driggers, V. R. Fantin, H. G. Jang, S. Jin, M. C. Keenan, K. M. Marks, R. M. Prins, P. S. Ward, K. E. Yen, L. M. Liau, J. D. Rabinowitz, L. C. Cantley, C. B. Thompson, M. G. Vander Heiden, S. M. Su, Cancer-associated IDH1 mutations produce 2-hydroxyglutarate. Nature 462, 739–44 (2009).

3. M. E. Figueroa, O. Abdel-Wahab, C. Lu, P. S. Ward, J. Patel, A. Shih, Y. Li, N. Bhagwat, A. Vasanthakumar, H. F. Fernandez, M. S. Tallman, Z. Sun, K. Wolniak, J. K. Peeters, W. Liu, S. E. Choe, V. R. Fantin, E. Paietta, B. Lowenberg, J. D. Licht, L. A. Godley, R. Delwel, P. J. Valk, C. B. Thompson, R. L. Levine, A. Melnick, Leukemic IDH1 and IDH2 mutations result in a hypermethylation phenotype, disrupt TET2 function, and impair hematopoietic differentiation. Cancer Cell 18, 553–67 (2010).

4. H. Noushmehr, D. J. Weisenberger, K. Diefes, H. S. Phillips, K. Pujara, B. P. Berman, F. Pan, C. E. Pelloski, E. P. Sulman, K. P. Bhat, R. G. Verhaak, K. A. Hoadley, D. N. Hayes, C. M. Perou, H. K. Schmidt, L. Ding, R. K. Wilson, D. Van Den Berg, H. Shen, H. Bengtsson, P. Neuvial, L. M. Cope, J. Buckley, J. G. Herman, S. B. Baylin, P. W. Laird, K. Aldape, Identification of a CpG island methylator phenotype that defines a distinct subgroup of glioma. Cancer Cell 17, 510–22 (2010).

5. F. Farshidfar, S. Zheng, M.-C. Gingras, Y. Newton, J. Shih, A. G. Robertson, T. Hinoue, K. A. Hoadley, E. A. Gibb, J. Roszik, K. R. Covington, C.-C. Wu, E. Shinbrot, N. Stransky, A. Hegde, J. D. Yang, E. Reznik, S. Sadeghi, C. S. Pedamallu, A. I. Ojesina, J. M. Hess, J. T. Auman, S. K. Rhie, R. Bowlby, M. J. Borad, Cancer Genome Atlas Network, A. X. Zhu, J. M. Stuart, C. Sander, R. Akbani, A. D. Cherniack, V. Deshpande, T. Mounajjed, W. C. Foo, M. S. Torbenson, D. E. Kleiner, P. W. Laird, D. A. Wheeler, A. J. McRee, O. F. Bathe, J. B. Andersen, N. Bardeesy, L. R. Roberts, L. N. Kwong, Integrative Genomic Analysis of Cholangiocarcinoma Identifies Distinct IDH-Mutant Molecular Profiles. Cell Rep 19, 2878– 2880 (2017).

6. W. A. Flavahan, Y. Drier, B. B. Liau, S. M. Gillespie, A. S. Venteicher, A. O. Stemmer-Rachamimov, M. L. Suva, B. E. Bernstein, Insulator dysfunction and oncogene activation in IDH mutant gliomas. Nature 529, 110–4 (2016).

7. W. A. Flavahan, Y. Drier, S. E. Johnstone, M. L. Hemming, D. R. Tarjan, E. Hegazi, S. J. Shareef, N. M. Javed, C. P. Raut, B. K. Eschle, P. C. Gokhale, J. L. Hornick, E. T. Sicinska, G. D. Demetri, B. E. Bernstein, Altered chromosomal topology drives oncogenic programs in SDH-deficient GISTs. Nature 575, 229–233 (2019).

8. G. J. Rahme, N. M. Javed, K. L. Puorro, S. Xin, V. Hovestadt, S. E. Johnstone, B. E. Bernstein, Modeling epigenetic lesions that cause gliomas. Cell 186, 3674–3685.e14 (2023).

9. M.-J. Wu, H. Kondo, A. V. Kammula, L. Shi, Y. Xiao, S. Dhiab, Q. Xu, C. J. Slater, O. I. Avila, J. Merritt, H. Kato, P. Kattel, J. Sussman, I. Gritti, J. Eccleston, Y. Sun, H. M. Cho, K. Olander, T. Katsuda, D. D. Shi, M. R. Savani, B. C. Smith, J. M. Cleary, R. Mostoslavsky, V. Vijay, Y. Kitagawa, H. Wakimoto, R. W. Jenkins, K. B. Yates, J. Paik, A. Tassinari, D. H. Saatcioglu, A. E. Tron, W. Haas, D. Cahill, S. K. McBrayer, R. T. Manguso, N. Bardeesy, Mutant IDH1 inhibition induces dsDNA sensing to activate tumor immunity. Science 385, eadl6173 (2024).

10. S. K. Saha, C. A. Parachoniak, K. S. Ghanta, J. Fitamant, K. N. Ross, M. S. Najem, S. Gurumurthy, E. A. Akbay, D. Sia, H. Cornella, O. Miltiadous, C. Walesky, V. Deshpande, A. X. Zhu, A. F. Hezel, K. E. Yen, K. S. Straley, J. Travins, J. Popovici-Muller, C. Gliser, C. R. Ferrone, U. Apte, J. M. Llovet, K.-K. Wong, S. Ramaswamy, N. Bardeesy, Mutant IDH inhibits HNF-4α to block hepatocyte differentiation and promote biliary cancer. Nature 513, 110–114 (2014).

11. M. Sasaki, C. B. Knobbe, J. C. Munger, E. F. Lind, D. Brenner, A. Brustle, I. S. Harris, R. Holmes, A. Wakeham, J. Haight, A. You-Ten, W. Y. Li, S. Schalm, S. M. Su, C. Virtanen, G. Reifenberger, P. S. Ohashi, D. L. Barber, M. E. Figueroa, A. Melnick, J. C. Zuniga-Pflucker, T. W. Mak, IDH1(R132H) mutation increases murine haematopoietic progenitors and alters epigenetics. Nature, doi: nature11323 [pii] 10.1038/nature11323 (2012).

12. F. Wang, J. Travins, B. DeLaBarre, V. Penard-Lacronique, S. Schalm, E. Hansen, K. Straley, A. Kernytsky, W. Liu, C. Gliser, H. Yang, S. Gross, E. Artin, V. Saada, E. Mylonas, C. Quivoron, J. Popovici-Muller, J. O. Saunders, F. G. Salituro, S. Yan, S. Murray, W. Wei, Y. Gao, L. Dang, M. Dorsch, S. Agresta, D. P. Schenkein, S. A. Biller, S. M. Su, S. de Botton, K. E. Yen, Targeted inhibition of mutant IDH2 in leukemia cells induces cellular differentiation. Science 340, 622–6 (2013).

13. A. Kernytsky, F. Wang, E. Hansen, S. Schalm, K. Straley, C. Gliser, H. Yang, J. Travins, S. Murray, M. Dorsch, S. Agresta, D. P. Schenkein, S. A. Biller, S. M. Su, W. Liu, K. E. Yen, IDH2 mutation-induced histone and DNA hypermethylation is progressively reversed by small-molecule inhibition. Blood 125, 296–303 (2015).

14. E. M. Stein, C. D. DiNardo, D. A. Pollyea, A. T. Fathi, G. J. Roboz, J. K. Altman, R. M. Stone, D. J. DeAngelo, R. L. Levine, I. W. Flinn, H. M. Kantarjian, R. Collins, M. R. Patel, A. E. Frankel, A. Stein, M. A. Sekeres, R. T. Swords, B. C. Medeiros, C. Willekens, P. Vyas, A. Tosolini, Q. Xu, R. D. Knight, K. E. Yen, S. Agresta, S. de Botton, M. S. Tallman, Enasidenib in mutant IDH2 relapsed or refractory acute myeloid leukemia. Blood 130, 722– 731 (2017).

15. C. D. DiNardo, E. M. Stein, S. de Botton, G. J. Roboz, J. K. Altman, A. S. Mims, R. Swords, R. H. Collins, G. N. Mannis, D. A. Pollyea, W. Donnellan, A. T. Fathi, A. Pigneux, H. P. Erba, G. T. Prince, A. S. Stein, G. L. Uy, J. M. Foran, E. Traer, R. K. Stuart, M. L. Arellano, J. L. Slack, M. A. Sekeres, C. Willekens, S. Choe, H. Wang, V. Zhang, K. E. Yen, S. M. Kapsalis, H. Yang, D. Dai, B. Fan, M. Goldwasser, H. Liu, S. Agresta, B. Wu, E. C. Attar, M. S. Tallman, R. M. Stone, H. M. Kantarjian, Durable Remissions with Ivosidenib in IDH1-Mutated Relapsed or Refractory AML. N Engl J Med 378, 2386–2398 (2018).

16. T. Watanabe, S. Nobusawa, P. Kleihues, H. Ohgaki, IDH1 mutations are early events in the development of astrocytomas and oligodendrogliomas. Am J Pathol 174, 1149–53 (2009).

17. D. N. Louis, A. Perry, P. Wesseling, D. J. Brat, I. A. Cree, D. Figarella-Branger, C. Hawkins, H. K. Ng, S. M. Pfister, G. Reifenberger, R. Soffietti, A. von Deimling, D. W. Ellison, The 2021 WHO Classification of Tumors of the Central Nervous System: a summary. Neuro Oncol 23, 1231–1251 (2021).

18. I. Tirosh, A. S. Venteicher, C. Hebert, L. E. Escalante, A. P. Patel, K. Yizhak, J. M. Fisher, C. Rodman, C. Mount, M. G. Filbin, C. Neftel, N. Desai, J. Nyman, B. Izar, C. C. Luo, J. M. Francis, A. A. Patel, M. L. Onozato, N. Riggi, K. J. Livak, D. Gennert, R. Satija, B. V. Nahed, W. T. Curry, R. L. Martuza, R. Mylvaganam, A. J. Iafrate, M. P. Frosch, T. R. Golub, M. N. Rivera, G. Getz, O. Rozenblatt-Rosen, D. P. Cahill, M. Monje, B. E. Bernstein, D. N. Louis, A. Regev, M. L. Suvà, Single-cell RNA-seq supports a developmental hierarchy in human oligodendroglioma. Nature 539, 309–313 (2016).

19. A. S. Venteicher, I. Tirosh, C. Hebert, K. Yizhak, C. Neftel, M. G. Filbin, V. Hovestadt, L. E. Escalante, M. L. Shaw, C. Rodman, S. M. Gillespie, D. Dionne, C. C. Luo, H. Ravichandran, R. Mylvaganam, C. Mount, M. L. Onozato, B. V. Nahed, H. Wakimoto, W. T. Curry, A. J. Iafrate, M. N. Rivera, M. P. Frosch, T. R. Golub, P. K. Brastianos, G. Getz, A. P. Patel, M. Monje, D. P. Cahill, O. Rozenblatt-Rosen, D. N. Louis, B. E. Bernstein, A. Regev, M. L. Suvà, Decoupling genetics, lineages, and microenvironment in IDH-mutant gliomas by single-cell RNA-seq. Science 355, eaai8478 (2017).

20. K. C. Johnson, K. J. Anderson, E. T. Courtois, A. D. Gujar, F. P. Barthel, F. S. Varn, D. Luo, M. Seignon, E. Yi, H. Kim, M. R. H. Estecio, D. Zhao, M. Tang, N. E. Navin, R. Maurya, C. Y. Ngan, N. Verburg, P. C. de Witt Hamer, K. Bulsara, M. L. Samuels, S. Das, P. Robson, R. G. W. Verhaak, Single-cell multimodal glioma analyses identify epigenetic regulators of cellular plasticity and environmental stress response. Nat Genet 53, 1456–1468 (2021).

21. A. Spitzer, S. Gritsch, M. Nomura, A. Jucht, J. Fortin, R. Raviram, H. R. Weisman, L. N. Gonzalez Castro, N. Druck, R. Chanoch-Myers, J. J. Y. Lee, R. Mylvaganam, R. Lee Servis, J. M. Fung, C. K. Lee, H. Nagashima, J. J. Miller, I. Arrillaga-Romany, D. N. Louis, H. Wakimoto, W. Pisano, P. Y. Wen, T. W. Mak, M. Sanson, M. Touat, D. A. Landau, K. L. Ligon, D. P. Cahill, M. L. Suvà, I. Tirosh, Mutant IDH inhibitors induce lineage differentiation in IDH-mutant oligodendroglioma. Cancer Cell 42, 904–914.e9 (2024).

22. C. Sojka, H.-L. V. Wang, T. N. Bhatia, Y. Li, P. Chopra, A. Sing, A. Voss, A. King, F. Wang, K. Joseph, V. M. Ravi, J. Olson, K. Hoang, E. Nduom, V. G. Corces, B. Yao, S. A. Sloan, Mapping the developmental trajectory of human astrocytes reveals divergence in glioblastoma. Nat Cell Biol 27, 347–359 (2025).

23. S. Turcan, D. Rohle, A. Goenka, L. A. Walsh, F. Fang, E. Yilmaz, C. Campos, A. W. Fabius, C. Lu, P. S. Ward, C. B. Thompson, A. Kaufman, O. Guryanova, R. Levine, A. Heguy, A. Viale, L. G. Morris, J. T. Huse, I. K. Mellinghoff, T. A. Chan, IDH1 mutation is sufficient to establish the glioma hypermethylator phenotype. Nature 483, 479–83 (2012).

24. N. M. Amankulor, Y. Kim, S. Arora, J. Kargl, F. Szulzewsky, M. Hanke, D. H. Margineantu, A. Rao, H. Bolouri, J. Delrow, D. Hockenbery, A. M. Houghton, E. C. Holland, Mutant IDH1 regulates the tumor-associated immune system in gliomas. Genes Dev 31, 774–786 (2017).

25. F. J. Núñez, F. M. Mendez, P. Kadiyala, M. S. Alghamri, M. G. Savelieff, M. B. Garcia-Fabiani, S. Haase, C. Koschmann, A.-A. Calinescu, N. Kamran, M. Saxena, R. Patel, S. Carney, M. Z. Guo, M. Edwards, M. Ljungman, T. Qin, M. A. Sartor, R. Tagett, S. Venneti, J. Brosnan-Cashman, A. Meeker, V. Gorbunova, L. Zhao, D. M. Kremer, L. Zhang, C. A. Lyssiotis, L. Jones, C. J. Herting, J. L. Ross, D. Hambardzumyan, S. Hervey-Jumper, M. E. Figueroa, P. R. Lowenstein, M. G. Castro, IDH1-R132H acts as a tumor suppressor in glioma via epigenetic up-regulation of the DNA damage response. Sci Transl Med 11, eaaq1427 (2019).

26. M. Sasaki, C. B. Knobbe, M. Itsumi, A. J. Elia, I. S. Harris, Chio, R. A. Cairns, S. McCracken, A. Wakeham, J. Haight, A. Y. Ten, B. Snow, T. Ueda, S. Inoue, K. Yamamoto, M. Ko, A. Rao, K. E. Yen, S. M. Su, T. W. Mak, D-2-hydroxyglutarate produced by mutant IDH1 perturbs collagen maturation and basement membrane function. Genes Dev 26, 2038– 49 (2012).

27. J. R. Adams, K. Xu, J. C. Liu, N. M. Agamez, A. J. Loch, R. G. Wong, W. Wang, K. L. Wright, T. F. Lane, E. Zacksenhaus, S. E. Egan, Cooperation between Pik3ca and p53 mutations in mouse mammary tumor formation. Cancer Res 71, 2706–17 (2011).

28. D. D. Shi, M. R. Savani, M. M. Levitt, A. C. Wang, J. E. Endress, C. E. Bird, J. Buehler, S. A. Stopka, M. S. Regan, Y.-F. Lin, V. T. Puliyappadamba, W. Gao, J. Khanal, L. Evans, J. H. Lee, L. Guo, Y. Xiao, M. Xu, B. Huang, R. B. Jennings, D. M. Bonal, M. S. Martin-Sandoval, T. Dang, L. C. Gattie, A. B. Cameron, S. Lee, J. M. Asara, H. I. Kornblum, T. W. Mak, R. E. Looper, Q.-D. Nguyen, S. Signoretti, S. Gradl, A. Sutter, M. Jeffers, A. Janzer, M. A. Lehrman, L. G. Zacharias, T. P. Mathews, J.-A. Losman, T. E. Richardson, D. P. Cahill, R. J. DeBerardinis, K. L. Ligon, L. Xu, P. Ly, N. Y. R. Agar, K. G. Abdullah, I. S. Harris, W. G. Kaelin, S. K. McBrayer, De novo pyrimidine synthesis is a targetable vulnerability in IDH mutant glioma. Cancer Cell 40, 939–956.e16 (2022).

29. K. Aoki, H. Nakamura, H. Suzuki, K. Matsuo, K. Kataoka, T. Shimamura, K. Motomura, F. Ohka, S. Shiina, T. Yamamoto, Y. Nagata, T. Yoshizato, M. Mizoguchi, T. Abe, Y. Momii, Y. Muragaki, R. Watanabe, I. Ito, M. Sanada, H. Yajima, N. Morita, I. Takeuchi, S. Miyano, T. Wakabayashi, S. Ogawa, A. Natsume, Prognostic relevance of genetic alterations in diffuse lower-grade gliomas. Neuro Oncol 20, 66–77 (2018).

30. J. Ganz, E. A. Maury, B. Becerra, S. Bizzotto, R. N. Doan, C. J. Kenny, T. Shin, J. Kim, Z. Zhou, K. L. Ligon, E. A. Lee, C. A. Walsh, Rates and Patterns of Clonal Oncogenic Mutations in the Normal Human Brain. Cancer Discov 12, 172–185 (2022).

31. D. D. Shi, J. H. Lee, A. C. Wang, J. Khanal, W. Gao, W. G. Kaelin, S. K. McBrayer, Protocol to establish a genetically engineered mouse model of IDH1-mutant astrocytoma. STAR Protoc 4, 102281 (2023).

32. R. Gao, S. Bai, Y. C. Henderson, Y. Lin, A. Schalck, Y. Yan, T. Kumar, M. Hu, E. Sei, A. Davis, F. Wang, S. F. Shaitelman, J. R. Wang, K. Chen, S. Moulder, S. Y. Lai, N. E. Navin, Delineating copy number and clonal substructure in human tumors from single-cell transcriptomes. Nat Biotechnol 39, 599–608 (2021).

33. Cancer Genome Atlas Research Network, D. J. Brat, R. G. W. Verhaak, K. D. Aldape, W. K. A. Yung, S. R. Salama, L. A. D. Cooper, E. Rheinbay, C. R. Miller, M. Vitucci, O. Morozova, A. G. Robertson, H. Noushmehr, P. W. Laird, A. D. Cherniack, R. Akbani, J. T. Huse, G. Ciriello, L. M. Poisson, J. S. Barnholtz-Sloan, M. S. Berger, C. Brennan, R. R. Colen, H. Colman, A. E. Flanders, C. Giannini, M. Grifford, A. Iavarone, R. Jain, I. Joseph, J. Kim, K. Kasaian, T. Mikkelsen, B. A. Murray, B. P. O’Neill, L. Pachter, D. W. Parsons, C. Sougnez, E. P. Sulman, S. R. Vandenberg, E. G. Van Meir, A. von Deimling, H. Zhang, D. Crain, K. Lau, D. Mallery, S. Morris, J. Paulauskis, R. Penny, T. Shelton, M. Sherman, P. Yena, A. Black, J. Bowen, K. Dicostanzo, J. Gastier-Foster, K. M. Leraas, T. M. Lichtenberg, C. R. Pierson, N. C. Ramirez, C. Taylor, S. Weaver, L. Wise, E. Zmuda, T. Davidsen, J. A. Demchok, G. Eley, M. L. Ferguson, C. M. Hutter, K. R. Mills Shaw, B. A. Ozenberger, M. Sheth, H. J. Sofia, R. Tarnuzzer, Z. Wang, L. Yang, J. C. Zenklusen, B. Ayala, J. Baboud, S. Chudamani, M. A. Jensen, J. Liu, T. Pihl, R. Raman, Y. Wan, Y. Wu, A. Ally, J. T. Auman, M. Balasundaram, S. Balu, S. B. Baylin, R. Beroukhim, M. S. Bootwalla, R. Bowlby, C. A. Bristow, D. Brooks, Y. Butterfield, R. Carlsen, S. Carter, L. Chin, A. Chu, E. Chuah, K. Cibulskis, A. Clarke, S. G. Coetzee, N. Dhalla, T. Fennell, S. Fisher, S. Gabriel, G. Getz, R. Gibbs, R. Guin, A. Hadjipanayis, D. N. Hayes, T. Hinoue, K. Hoadley, R. A. Holt, A. P. Hoyle, S. R. Jefferys, S. Jones, C. D. Jones, R. Kucherlapati, P. H. Lai, E. Lander, S. Lee, L. Lichtenstein, Y. Ma, D. T. Maglinte, H. S. Mahadeshwar, M. A. Marra, M. Mayo, S. Meng, M. L. Meyerson, P. A. Mieczkowski, R. A. Moore, L. E. Mose, A. J. Mungall, A. Pantazi, M. Parfenov, P. J. Park, J. S. Parker, C. M. Perou, A. Protopopov, X. Ren, J. Roach, T. S. Sabedot, J. Schein, S. E. Schumacher, J. G. Seidman, S. Seth, H. Shen, J. V. Simons, P. Sipahimalani, M. G. Soloway, X. Song, H. Sun, B. Tabak, A. Tam, D. Tan, J. Tang, N. Thiessen, T. Triche, D. J. Van Den Berg, U. Veluvolu, S. Waring, D. J. Weisenberger, M. D. Wilkerson, T. Wong, J. Wu, L. Xi, A. W. Xu, L. Yang, T. I. Zack, J. Zhang, B. A. Aksoy, H. Arachchi, C. Benz, B. Bernard, D. Carlin, J. Cho, D. DiCara, S. Frazer, G. N. Fuller, J. Gao, N. Gehlenborg, D. Haussler, D. I. Heiman, L. Iype, A. Jacobsen, Z. Ju, S. Katzman, H. Kim, T. Knijnenburg, R. B. Kreisberg, M. S. Lawrence, W. Lee, K. Leinonen, P. Lin, S. Ling, W. Liu, Y. Liu, Y. Liu, Y. Lu, G. Mills, S. Ng, M. S. Noble, E. Paull, A. Rao, S. Reynolds, G. Saksena, Z. Sanborn, C. Sander, N. Schultz, Y. Senbabaoglu, R. Shen, I. Shmulevich, R. Sinha, J. Stuart, S. O. Sumer, Y. Sun, N. Tasman, B. S. Taylor, D. Voet, N. Weinhold, J. N. Weinstein, D. Yang, K. Yoshihara, S. Zheng, W. Zhang, L. Zou, T. Abel, S. Sadeghi, M. L. Cohen, J. Eschbacher, E. M. Hattab, A. Raghunathan, M. J. Schniederjan, D. Aziz, G. Barnett, W. Barrett, D. D. Bigner, L. Boice, C. Brewer, C. Calatozzolo, B. Campos, C. G. Carlotti, T. A. Chan, L. Cuppini, E. Curley, S. Cuzzubbo, K. Devine, F. DiMeco, R. Duell, J. B. Elder, A. Fehrenbach, G. Finocchiaro, W. Friedman, J. Fulop, J. Gardner, B. Hermes, C. Herold-Mende, C. Jungk, A. Kendler, N. L. Lehman, E. Lipp, O. Liu, R. Mandt, M. McGraw, R. Mclendon, C. McPherson, L. Neder, P. Nguyen, A. Noss, R. Nunziata, Q. T. Ostrom, C. Palmer, A. Perin, B. Pollo, A. Potapov, O. Potapova, W. K. Rathmell, D. Rotin, L. Scarpace, C. Schilero, K. Senecal, K. Shimmel, V. Shurkhay, S. Sifri, R. Singh, A. E. Sloan, K. Smolenski, S. M. Staugaitis, R. Steele, L. Thorne, D. P. C. Tirapelli, A. Unterberg, M. Vallurupalli, Y. Wang, R. Warnick, F. Williams, Y. Wolinsky, S. Bell, M. Rosenberg, C. Stewart, F. Huang, J. L. Grimsby, A. J. Radenbaugh, J. Zhang, Comprehensive, Integrative Genomic Analysis of Diffuse Lower-Grade Gliomas. N Engl J Med 372, 2481–2498 (2015).

34. G.-L. Ming, H. Song, Adult neurogenesis in the mammalian brain: significant answers and significant questions. Neuron 70, 687–702 (2011).

35. J. Wu, L. N. Gonzalez Castro, S. Battaglia, C. A. El Farran, J. P. D’Antonio, T. E. Miller, M. L. Suvà, B. E. Bernstein, Evolving cell states and oncogenic drivers during the progression of IDH-mutant gliomas. Nat Cancer 6, 145–157 (2025).

36. I. K. Mellinghoff, M. Lu, P. Y. Wen, J. W. Taylor, E. A. Maher, I. Arrillaga-Romany, K. B. Peters, B. M. Ellingson, M. K. Rosenblum, S. Chun, K. Le, A. Tassinari, S. Choe, Y. Toubouti, S. Schoenfeld, S. S. Pandya, I. Hassan, L. Steelman, J. L. Clarke, T. F. Cloughesy, Vorasidenib and ivosidenib in IDH1-mutant low-grade glioma: a randomized, perioperative phase 1 trial. Nat Med 29, 615–622 (2023).

37. K. Roychoudhury, J. Salomone, S. Qin, B. Cain, M. Adam, S. S. Potter, M. Nakafuku, B. Gebelein, K. Campbell, Physical interactions between Gsx2 and Ascl1 balance progenitor expansion versus neurogenesis in the mouse lateral ganglionic eminence. Development 147, dev185348 (2020).

38. H. Chapman, R. R. Waclaw, Z. Pei, M. Nakafuku, K. Campbell, The homeobox gene Gsx2 controls the timing of oligodendroglial fate specification in mouse lateral ganglionic eminence progenitors. Development 140, 2289–2298 (2013).

39. M. A. Petryniak, G. B. Potter, D. H. Rowitch, J. L. R. Rubenstein, Dlx1 and Dlx2 control neuronal versus oligodendroglial cell fate acquisition in the developing forebrain. Neuron 55, 417–433 (2007).

40. H. Chapman, A. Riesenberg, L. A. Ehrman, V. Kohli, D. Nardini, M. Nakafuku, K. Campbell, R. R. Waclaw, Gsx transcription factors control neuronal versus glial specification in ventricular zone progenitors of the mouse lateral ganglionic eminence. Dev Biol 442, 115–126 (2018).

41. Z. Pei, B. Wang, G. Chen, M. Nagao, M. Nakafuku, K. Campbell, Homeobox genes Gsx1 and Gsx2 differentially regulate telencephalic progenitor maturation. Proc Natl Acad Sci U S A 108, 1675–1680 (2011).

42. E. Letouzé, C. Martinelli, C. Loriot, N. Burnichon, N. Abermil, C. Ottolenghi, M. Janin, M. Menara, A. T. Nguyen, P. Benit, A. Buffet, C. Marcaillou, J. Bertherat, L. Amar, P. Rustin, A. De Reyniès, A.-P. Gimenez-Roqueplo, J. Favier, SDH mutations establish a hypermethylator phenotype in paraganglioma. Cancer Cell 23, 739–752 (2013).

43. Cancer Genome Atlas Research Network, W. M. Linehan, P. T. Spellman, C. J. Ricketts, C. J. Creighton, S. S. Fei, C. Davis, D. A. Wheeler, B. A. Murray, L. Schmidt, C. D. Vocke, M. Peto, A. A. M. Al Mamun, E. Shinbrot, A. Sethi, S. Brooks, W. K. Rathmell, A. N. Brooks, K. A. Hoadley, A. G. Robertson, D. Brooks, R. Bowlby, S. Sadeghi, H. Shen, D. J. Weisenberger, M. Bootwalla, S. B. Baylin, P. W. Laird, A. D. Cherniack, G. Saksena, S. Haake, J. Li, H. Liang, Y. Lu, G. B. Mills, R. Akbani, M. D. M. Leiserson, B. J. Raphael, P. Anur, D. Bottaro, L. Albiges, N. Barnabas, T. K. Choueiri, B. Czerniak, A. K. Godwin, A. A. Hakimi, T. H. Ho, J. Hsieh, M. Ittmann, W. Y. Kim, B. Krishnan, M. J. Merino, K. R. Mills Shaw, V. E. Reuter, E. Reznik, C. S. Shelley, B. Shuch, S. Signoretti, R. Srinivasan, P. Tamboli, G. Thomas, S. Tickoo, K. Burnett, D. Crain, J. Gardner, K. Lau, D. Mallery, S. Morris, J. D. Paulauskis, R. J. Penny, C. Shelton, W. T. Shelton, M. Sherman, E. Thompson, P. Yena, M. T. Avedon, J. Bowen, J. M. Gastier-Foster, M. Gerken, K. M. Leraas, T. M. Lichtenberg, N. C. Ramirez, T. Santos, L. Wise, E. Zmuda, J. A. Demchok, I. Felau, C. M. Hutter, M. Sheth, H. J. Sofia, R. Tarnuzzer, Z. Wang, L. Yang, J. C. Zenklusen, J. Zhang, B. Ayala, J. Baboud, S. Chudamani, J. Liu, L. Lolla, R. Naresh, T. Pihl, Q. Sun, Y. Wan, Y. Wu, A. Ally, M. Balasundaram, S. Balu, R. Beroukhim, T. Bodenheimer, C. Buhay, Y. S. N. Butterfield, R. Carlsen, S. L. Carter, H. Chao, E. Chuah, A. Clarke, K. R. Covington, M. Dahdouli, N. Dewal, N. Dhalla, H. V. Doddapaneni, J. A. Drummond, S. B. Gabriel, R. A. Gibbs, R. Guin, W. Hale, A. Hawes, D. N. Hayes, R. A. Holt, A. P. Hoyle, S. R. Jefferys, S. J. M. Jones, C. D. Jones, D. Kalra, C. Kovar, L. Lewis, J. Li, Y. Ma, M. A. Marra, M. Mayo, S. Meng, M. Meyerson, P. A. Mieczkowski, R. A. Moore, D. Morton, L. E. Mose, A. J. Mungall, D. Muzny, J. S. Parker, C. M. Perou, J. Roach, J. E. Schein, S. E. Schumacher, Y. Shi, J. V. Simons, P. Sipahimalani, T. Skelly, M. G. Soloway, C. Sougnez, A. Tam, D. Tan, N. Thiessen, U. Veluvolu, M. Wang, M. D. Wilkerson, T. Wong, J. Wu, L. Xi, J. Zhou, J. Bedford, F. Chen, Y. Fu, M. Gerstein, D. Haussler, K. Kasaian, P. Lai, S. Ling, A. Radenbaugh, D. Van Den Berg, J. N. Weinstein, J. Zhu, M. Albert, I. Alexopoulou, J. J. Andersen, J. T. Auman, J. Bartlett, S. Bastacky, J. Bergsten, M. L. Blute, L. Boice, R. J. Bollag, J. Boyd, E. Castle, Y.-B. Chen, J. C. Cheville, E. Curley, B. Davies, A. DeVolk, R. Dhir, L. Dike, J. Eckman, J. Engel, J. Harr, R. Hrebinko, M. Huang, L. Huelsenbeck-Dill, M. Iacocca, B. Jacobs, M. Lobis, J. K. Maranchie, S. McMeekin, J. Myers, J. Nelson, J. Parfitt, A. Parwani, N. Petrelli, B. Rabeno, S. Roy, A. L. Salner, J. Slaton, M. Stanton, R. H. Thompson, L. Thorne, K. Tucker, P. M. Weinberger, C. Winemiller, L. A. Zach, R. Zuna, Comprehensive Molecular Characterization of Papillary Renal-Cell Carcinoma. N Engl J Med 374, 135–145 (2016).

44. A. López-Juárez, J. Howard, K. Ullom, L. Howard, A. Grande, A. Pardo, R. Waclaw, Y.-Y. Sun, D. Yang, C.-Y. Kuan, K. Campbell, M. Nakafuku, Gsx2 controls region-specific activation of neural stem cells and injury-induced neurogenesis in the adult subventricular zone. Genes Dev 27, 1272–1287 (2013).

45. C. C. L. Chen, S. Deshmukh, S. Jessa, D. Hadjadj, V. Lisi, A. F. Andrade, D. Faury, W. Jawhar, R. Dali, H. Suzuki, M. Pathania, D. A, F. Dubois, E. Woodward, S. Hébert, M. Coutelier, J. Karamchandani, S. Albrecht, S. Brandner, N. De Jay, T. Gayden, A. Bajic, A. S. Harutyunyan, D. M. Marchione, L. G. Mikael, N. Juretic, M. Zeinieh, C. Russo, N. Maestro, A. V. Bassenden, P. Hauser, J. Virga, L. Bognar, A. Klekner, M. Zapotocky, A. Vicha, L. Krskova, K. Vanova, J. Zamecnik, D. Sumerauer, P. G. Ekert, D. S. Ziegler, B. Ellezam, M. G. Filbin, M. Blanchette, J. R. Hansford, D.-A. Khuong-Quang, A. M. Berghuis, A. G. Weil, A. A. Garcia, L. Garzia, S. C. Mack, R. Beroukhim, K. L. Ligon, M. D. Taylor, P. Bandopadhayay, C. Kramm, S. M. Pfister, A. Korshunov, D. Sturm, D. T. W. Jones, P. Salomoni, C. L. Kleinman, N. Jabado, Histone H3.3G34-Mutant Interneuron Progenitors Co-opt PDGFRA for Gliomagenesis. Cell 183, 1617–1633.e22 (2020).

46. Y. Wu, M. Fletcher, Z. Gu, Q. Wang, B. Costa, A. Bertoni, K.-H. Man, M. Schlotter, J. Felsberg, J. Mangei, M. Barbus, A.-C. Gaupel, W. Wang, T. Weiss, R. Eils, M. Weller, H. Liu, G. Reifenberger, A. Korshunov, P. Angel, P. Lichter, C. Herrmann, B. Radlwimmer, Glioblastoma epigenome profiling identifies SOX10 as a master regulator of molecular tumour subtype. Nat Commun 11, 6434 (2020).

47. R. S. Guerra-Resendez, S. L. Lydon, A. J. Ma, G. C. Bedford, D. R. Reed, S. Kim, E. R. Teran, T. Nishiguchi, M. Escobar, A. R. DiNardo, I. B. Hilton, Characterization of Rationally Designed CRISPR/Cas9-Based DNA Methyltransferases with Distinct Methyltransferase and Gene Silencing Activities in Human Cell Lines and Primary Human T Cells. ACS Synth. Biol. 14, 384–397 (2025).

48. C. Bardella, O. Al-Dalahmah, D. Krell, P. Brazauskas, K. Al-Qahtani, M. Tomkova, J. Adam, S. Serres, H. Lockstone, L. Freeman-Mills, I. Pfeffer, N. Sibson, R. Goldin, B. Schuster-Boeckler, P. J. Pollard, T. Soga, J. S. McCullagh, C. J. Schofield, P. Mulholland, O. Ansorge, S. Kriaucionis, P. J. Ratcliffe, F. G. Szele, I. Tomlinson, Expression of Idh1(R132H) in the Murine Subventricular Zone Stem Cell Niche Recapitulates Features of Early Gliomagenesis. Cancer Cell 30, 578–594 (2016).

49. H. Zong, L. F. Parada, S. J. Baker, Cell of origin for malignant gliomas and its implication in therapeutic development. Cold Spring Harb Perspect Biol 7, a020610 (2015).

50. C. Liu, J. C. Sage, M. R. Miller, R. G. W. Verhaak, S. Hippenmeyer, H. Vogel, O. Foreman, R. T. Bronson, A. Nishiyama, L. Luo, H. Zong, Mosaic analysis with double markers reveals tumor cell of origin in glioma. Cell 146, 209–221 (2011).

51. J. H. Lee, J. E. Lee, J. Y. Kahng, S. H. Kim, J. S. Park, S. J. Yoon, J.-Y. Um, W. K. Kim, J.- K. Lee, J. Park, E. H. Kim, J.-H. Lee, J.-H. Lee, W.-S. Chung, Y. S. Ju, S.-H. Park, J. H. Chang, S.-G. Kang, J. H. Lee, Human glioblastoma arises from subventricular zone cells with low-level driver mutations. Nature 560, 243–247 (2018).

52. R. G. Verhaak, K. A. Hoadley, E. Purdom, V. Wang, Y. Qi, M. D. Wilkerson, C. R. Miller, L. Ding, T. Golub, J. P. Mesirov, G. Alexe, M. Lawrence, M. O’Kelly, P. Tamayo, B. A. Weir, S. Gabriel, W. Winckler, S. Gupta, L. Jakkula, H. S. Feiler, J. G. Hodgson, C. D. James, J. N. Sarkaria, C. Brennan, A. Kahn, P. T. Spellman, R. K. Wilson, T. P. Speed, J. W. Gray, M. Meyerson, G. Getz, C. M. Perou, D. N. Hayes, Integrated genomic analysis identifies clinically relevant subtypes of glioblastoma characterized by abnormalities in PDGFRA, IDH1, EGFR, and NF1. Cancer Cell 17, 98–110 (2010).

53. J. W. Park, J. Kwak, K.-W. Kim, S. Jung, C. H. Nam, H. J. Kim, S. M. Lee, J.-H. Park, J. Yoo, J.-K. Shim, C. Kim, S. Ahn, S. Pusch, A. Von Deimling, J. H. Chang, S. H. Kim, Y. S. Ju, S.-G. Kang, J. H. Lee, IDH-mutant gliomas arise from glial progenitor cells harboring the initial driver mutation. Cold Spring Harbor Laboratory [Preprint] (2024). 10.1101/2024.10.17.618976.

54. I. Liu, G. Alencastro Veiga Cruzeiro, L. Bjerke, R. F. Rogers, Y. Grabovska, A. Beck, A. Mackay, T. Barron, O. A. Hack, M. A. Quezada, V. Molinari, M. L. Shaw, M. Perez-Somarriba, S. Temelso, F. Raynaud, R. Ruddle, E. Panditharatna, B. Englinger, H. M. Mire, L. Jiang, A. Nascimento, J. LaBelle, R. Haase, J. Rozowsky, S. Neyazi, A.-C. Baumgartner, S. Castellani, S. E. Hoffman, A. Cameron, M. Morrow, Q.-D. Nguyen, G. Pericoli, S. Madlener, L. Mayr, C. Dorfer, R. Geyeregger, C. Rota, G. Ricken, K. L. Ligon, S. Alexandrescu, R. T. Cartaxo, B. Lau, S. Uphadhyaya, C. Koschmann, E. Braun, M. Danan-Gotthold, L. Hu, K. Siletti, E. Sundström, R. Hodge, E. Lein, S. Agnihotri, D. D. Eisenstat, S. Stapleton, A. King, C. Bleil, A. Mastronuzzi, K. A. Cole, A. J. Waanders, A. Montero Carcaboso, U. Schüller, D. Hargrave, M. Vinci, F. Carceller, C. Haberler, I. Slavc, S. Linnarsson, J. Gojo, M. Monje, C. Jones, M. G. Filbin, GABAergic neuronal lineage development determines clinically actionable targets in diffuse hemispheric glioma, H3G34-mutant. Cancer Cell, S1535-6108(24)00305–2 (2024).

55. W. Xu, H. Yang, Y. Liu, Y. Yang, P. Wang, S. H. Kim, S. Ito, C. Yang, M. T. Xiao, L. X. Liu, W. Q. Jiang, J. Liu, J. Y. Zhang, B. Wang, S. Frye, Y. Zhang, Y. H. Xu, Q. Y. Lei, K. L. Guan, S. M. Zhao, Y. Xiong, Oncometabolite 2-hydroxyglutarate is a competitive inhibitor of alpha-ketoglutarate-dependent dioxygenases. Cancer Cell 19, 17–30 (2011).

56. R. R. Waclaw, B. Wang, Z. Pei, L. A. Ehrman, K. Campbell, Distinct temporal requirements for the homeobox gene Gsx2 in specifying striatal and olfactory bulb neuronal fates. Neuron 63, 451–465 (2009).

57. D. W. Parsons, S. Jones, X. Zhang, J. C. Lin, R. J. Leary, P. Angenendt, P. Mankoo, H. Carter, I. M. Siu, G. L. Gallia, A. Olivi, R. McLendon, B. A. Rasheed, S. Keir, T. Nikolskaya, Y. Nikolsky, D. A. Busam, H. Tekleab, L. A. Diaz, J. Hartigan, D. R. Smith, R. L. Strausberg, S. K. Marie, S. M. Shinjo, H. Yan, G. J. Riggins, D. D. Bigner, R. Karchin, N. Papadopoulos, G. Parmigiani, B. Vogelstein, V. E. Velculescu, K. W. Kinzler, An integrated genomic analysis of human glioblastoma multiforme. Science 321, 1807–12 (2008).

58. P. Koivunen, S. Lee, C. G. Duncan, G. Lopez, G. Lu, S. Ramkissoon, J. A. Losman, P. Joensuu, U. Bergmann, S. Gross, J. Travins, S. Weiss, R. Looper, K. L. Ligon, R. G. Verhaak, H. Yan, W. G. Kaelin, Transformation by the (R)-enantiomer of 2-hydroxyglutarate linked to EGLN activation. Nature 483, 484–488 (2012).

59. B. Blouw, H. Song, T. Tihan, J. Bosze, N. Ferrara, H. P. Gerber, R. S. Johnson, G. Bergers, The hypoxic response of tumors is dependent on their microenvironment. Cancer Cell 4, 133–46 (2003).

60. I. K. Mellinghoff, M. J. van den Bent, D. T. Blumenthal, M. Touat, K. B. Peters, J. Clarke, J. Mendez, S. Yust-Katz, L. Welsh, W. P. Mason, F. Ducray, Y. Umemura, B. Nabors, M. Holdhoff, A. F. Hottinger, Y. Arakawa, J. M. Sepulveda, W. Wick, R. Soffietti, J. R. Perry, P. Giglio, M. de la Fuente, E. A. Maher, S. Schoenfeld, D. Zhao, S. S. Pandya, L. Steelman, I. Hassan, P. Y. Wen, T. F. Cloughesy, Vorasidenib in IDH1- or IDH2-Mutant Low-Grade Glioma. N Engl J Med 389, 589–601 (2023).

