## Supplemental materials for "Mutant IDH silences GSX2 to reprogram neural progenitor cell fate and promote gliomagenesis"

**The PDF file includes:**

Materials and Methods
Figs. S1 to S12
References 1 to 22

**Other Supplementary Materials for this manuscript include the following:**

Data S1 to S5

**Materials and Methods**

Experimental Model Details

Animals

All care and treatment of experimental animals were carried out in strict accordance with Good Animal Practice as defined by the US Office of Laboratory Animal Welfare and approved by the UT Southwestern Medical Center Institutional Animal Care and Use Committee (protocol 2019–102795). Animal welfare was assessed daily throughout treatment periods. Mice were housed together (2–5 mice per cage) and provided free access to standard diet and water. Strains used included: R26$^{\text{LSL-Cas9-EGFP}}$ (B6J.129(B6N)-Gt(ROSA)26Sor$^{\text{tm1(CAG-cas9*,-EGFP)Fezh}}$/J, Jackson stock #026175) (*1*), R26$^{\text{LSL-Pik3ca-H1047R}}$ (FVB.129S6-Gt(ROSA)26Sor$^{\text{tm1(Pik3ca*H1047R)Egan}}$/J, Jackson stock #016977) (*2*), Idh1$^{\text{tm1Mak}}$ (backcrossed to C57BL/6, provided by T. Mak, University of Toronto) (*3*, *4*).

To generate the experimental cohorts for the survival study, we first bred transgenic Idh1$^{\text{tm1Mak/WT}}$ mice with R26$^{\text{LSL-Cas9-EGFP+/+}}$ mice to produce Idh1$^{\text{tm1Mak/WT}}$; R26$^{\text{LSL-Cas9-EGFP+/-}}$ progeny, which were subsequently interbred to produce Idh1$^{\text{tm1Mak/WT}}$; R26$^{\text{LSL-Cas9-EGFP+/+}}$ mice. Idh1$^{\text{tm1Mak/WT}}$; R26$^{\text{LSL-Cas9-EGFP+/+}}$ mice were crossed with R26$^{\text{LSL-Pik3ca-H1047R+/+}}$ to produce ***PIC*** (R26$^{\text{LSL-Pik3ca-H1047R+/-}}$; Idh1$^{\text{tm1Mak/WT}}$; R26$^{\text{LSL-Cas9-EGFP+/–}}$) and ***PC*** (R26$^{\text{LSL-Pik3ca-H1047R+/-}}$; Idh1$^{\text{WT/WT}}$; R26$^{\text{LSL-Cas9-EGFP+/–}}$) mice. Separately, Idh1$^{\text{tm1Mak/WT}}$; R26$^{\text{LSL-Cas9-EGFP+/+}}$ mice were crossed with wild-type FVB mice to produce ***IC*** (Idh1$^{\text{tm1Mak/WT}}$ ; R26$^{\text{LSL-Cas9-EGFP+/–}}$) and ***C*** (Idh1$^{\text{WT/WT}}$; R26$^{\text{LSL-Cas9-EGFP+/–}}$) mice.

To generate the experimental cohorts to monitor the cellular and molecular dynamics of gliomagenesis, we first bred transgenic Idh1$^{\text{tm1Mak/WT}}$ mice with R26$^{\text{LSL-Cas9-EGFP+/+}}$ mice to produce Idh1$^{\text{tm1Mak/WT}}$; R26$^{\text{LSL-Cas9-EGFP+/-}}$ progeny, which were subsequently interbred to produce homozygous Idh1$^{\text{tm1Mak/tm1Mak}}$; R26$^{\text{LSL-Cas9-EGFP+/+}}$ mice. Idh1$^{\text{tm1Mak/tm1Mak}}$; R26$^{\text{LSL-Cas9-EGFP+/+}}$ mice were crossed with R26$^{\text{LSL-Pik3ca-H1047R+/+}}$ to produce ***PIC*** (R26$^{\text{LSL-Pik3ca-H1047R+/-}}$; Idh1$^{\text{tm1Mak/WT}}$; R26$^{\text{LSL-Cas9-EGFP+/–}}$) mice. R26$^{\text{LSL-Cas9-EGFP+/+}}$ mice were crossed with R26$^{\text{LSL-Pik3ca-H1047R+/+}}$ to produce ***PC*** (R26$^{\text{LSL-Pik3ca-H1047R+/-}}$; Idh1$^{\text{WT/WT}}$; R26$^{\text{LSL-Cas9-EGFP+/–}}$) mice. R26$^{\text{LSL-Cas9-EGFP+/+}}$ mice were crossed with wild-type FVB mice to produce ***C*** (Idh1$^{\text{WT/WT}}$; R26$^{\text{LSL-Cas9-EGFP+/–}}$) mice.

Genotyping was performed by Transnetyx. For intracranial AAV injections, P2 pups were anesthetized by hypothermia and immobilized using a stereotactic frame. AAV particles (1 µL, > $10^{13}$ GC/mL) were injected into the subventricular zone using a 5 µL syringe (Hamilton, 7803-04). The whole injection procedure, including hypothermia-induced anesthesia, was kept under 10 minutes. Pups were allowed to recover on a pre-warmed heating pad and returned to their mother once normal body temperature and skin color were restored. Tumor size and survival analyses were performed by researchers not blinded to genotypes or treatment arms. Mice were euthanized upon exhibiting neurological symptoms or becoming moribund.

Primary Cell Lines

Primary mouse GSCs or NSCs derived from GEM models were cultured in the presence of 5% $CO_2$, 5% $O_2$ at 37°C in NeuroCult Basal Medium (Mouse and Rat) with Proliferation Supplement (StemCell Technologies, 05702) supplemented with EGF (20 ng/mL, Goldbio, 1150-04-500), bFGF (20 ng/mL, Goldbio, 1140-02-500), heparin (2 µg/mL, StemCell Technologies, 07980), 1% penicillin/streptomycin (Thermo Scientific, 15140163), amphotericin B (250 ng/mL, Fisher Scientific, SV3007801), and Plasmocin (0.25 µg/mL, Fisher Scientific,

NC1484222). All cell lines were routinely evaluated for mycoplasma contamination and tested negative throughout the study.

Methods

AAV plasmid cloning and AAV production

The pAAV2-sgTrp53-sgAtrx-Nes-Cre AAV expression vector was generated by replacing the EFS promoter in our previously used pAAV2-sgTrp53-sgAtrx-EFS-Cre construct (*5*) with a nestin enhancer linked to a TK minimal promoter (hereafter referred to as the nestin promoter) (*6*). The original construct was linearized by inverse PCR to remove the EFS promoter, and the nestin promoter fragment was PCR-amplified from the nes374tk/lacZ plasmid (Addgene #47615) and inserted into the linearized AAV backbone using InFusion cloning (Takara Bio, 639650). The AAV particles to generate the GEM models were prepared by Charles River Laboratories. For the *Gsx2* ablation experiments, the AAV expression plasmid was linearized by PCR, a DNA fragment containing U6 promoter, a sgRNA targeting *Hipp11* safe harbor locus (CACATAAGTTTAGACGCATT) or *Gsx2* (GATCTCGATTCTCCGGAGTC) and sgRNA scaffold was synthesized by TWIST Bioscience, then cloned into the AAV plasmid using NEBuilder HiFi DNA Assembly Master Mix (NEB, E2621L). AAV production and titration was done as previously described (*7*) with minor modifications. Briefly, about 70% confluent HEK293AAVpro cells grown in DMEM supplemented with 10% FBS were triple transfected with pAAV expression plasmid, pAAV Rep-Cap DJ (Cell Biolabs, VPK-420-DJ), and pHelper (Cell Biolabs, 340202) plasmids using PEI max (Kyfora Bio, 24765100) in 20 15 cm flasks. At day 4 post-transfection, media and cells were collected and processed separately. AAVs in cells were extracted using AAV Extraction Solution (Fisher Scientific, NC1415836). AAV particles were precipitated from medium with polyethylene glycol (PEG) 8000 (Sigma-Aldrich, 89510-250G-F). The PEG-precipitated AAV was collected by centrifugation and combined with AAVs extracted from cell pellet. AAVs were further purified using discontinuous iodixanol gradient ultracentrifugation. Primers binding within the AAV2 ITRs (Forward ITR primer: GGAACCCCTAGTGATGGAGTT, reverse ITR primer: CGGCCTCAGTGAGCGA) were used to measure the virus titer with quantitative polymerase chain reaction (qPCR). Before releasing the viral DNA from the particles, all extra-viral DNA was removed by digestion with DNase I. Then, DNase I was inactivated by adding EDTA. The viral DNA was released via AAV capsid lysis by adding proteinase K. The extracted viral DNA and a serial dilution of the pAAV expression plasmid as a standard were measured using the Luna Universal qPCR Master Mix (NEB, M3003L) on a Bio-Rad CFX384 real-time system.

Mouse brain dissociation and FACS

Mice at specific ages were euthanized, cardiac perfusion with phosphate-buffered saline (PBS) was performed and whole brain was extracted and immediately dissociated to single cell suspension using the neural tissue dissociation kit (Miltenyi Biotec 130-092-628). Cells were stained with calcein violet (Invitrogen, C34858) and ethidium homodimer-1 (Invitrogen, E1169) for 15 mins and sorted on a FACS Aria II SORP (4 lasers) from BD Biosciences. GFP+ live cells (calcein violet +, ethidium homodimer-1 -) were isolated for downstream sequencing analyses.

Single nucleus multiome sample preparation

Nuclei isolation was performed for the sorted GFP+ cells per 10X Genomics protocol (CG000365). Single-nucleus RNA-seq and ATAC-seq libraries were generated using the 10X Genomics Chromium Controller following the manufacturer's protocol for the Chromium Next GEM Single Cell Multiome ATAC + Gene Expression Reagent Bundle (10X Genomics, PN-1000283). Generated libraries were quantified by Qubit DNA high sensitivity assay (Thermo Fisher Scientific, Q33231) and fragment size was analyzed by Agilent Tapestation high sensitivity D5000 tapes. Library sequencing was performed using an Illumina NextSeq 500 at the Children's Research Institute's Sequencing Facility at UT Southwestern Medical Center. Raw reads were generated and further quantified using 10X Genomics Cell Ranger ARC (v2.0.2) with the 10X Genomics pre-built reference genome refdata-cellranger-arc-mm10-2020-A-2.0.0 for alignment at the UT Southwestern BioHPC.

Single nucleus multiome data analysis

Downstream analyses were performed using the R packages Seurat (V4.3.0) (*8*) and Signac (V1.13.0) (*9*). For snATAC-seq data, cells were retained based on the following criteria: total counts between 1,000 and 150,000, nucleosome signal less than 2, and TSS enrichment greater than 1. For snRNA-seq data, cells were retained if they met the following thresholds: mitochondrial gene expression less than 20%, fewer than 8,000 detected genes, and total counts between 1,000 and 20,000-50,000 (sample-specific upper limit). For integration and normalization, snRNA-seq and snATAC-seq datasets were processed separately. snATAC-seq samples were integrated using the "FindIntegrationAnchors" and "IntegrateEmbeddings" functions from Seurat and Signac packages. Peaks were annotated using the Bioconductor annotation package EnsDb.Mmusculus.v79. snRNA-seq samples were integrated using the reciprocal PCA method. The two modalities were then jointly analyzed using weighted nearest neighbor analysis via the "FindMultiModalNeighbors" function. Clusters were identified using "FindClusters" at a resolution of 0.3. Cluster marker genes were identified using the "FindAllMarkers" function, and cell types were annotated based on marker gene expression patterns. Ploidy status was inferred using the R package CopyKAT (*10*). The UMI count matrix from snRNA-seq was exported from the Seurat object and used as input for CopyKAT. Cells inferred to be aneuploid by CopyKAT were annotated as malignant, while diploid cells were annotated as non-transformed. Ploidy inference was consistent regardless of whether cells from C mice were specified as a diploid reference population or no reference diploid population was specified. For the pesudotime analysis, the Seurat object containing the undifferentiated neural cell and interneuron/neuroblast populations was converted to a Monocle object using the "as.cell_data_set" function from the R package SeuratWrappers, and the trajectory analysis was done by using Monocle3 (*11*).

Malignant cell states scoring

Scoring human and murine IDH-mutant astrocytoma cells was done similarly as described previously (*12*, *13*). Briefly, human and murine malignant cells (aneuploid cells defined by CopyKAT as above) were subsetted, and scored for the AC-like, OC-like, G1S, G2M and stemness programs (*13*) using the sigScores function from the R package scalop (https://github.com/jlaffy/scalop). Single cell RNA-seq data for human IDH-mutant astrocytomas were from Venteicher et al. 2017 (*14*). The cell lineage and stemness score of each cell were calculated as in Tirosh et al. 2016 (*12*). Cycling cells were defined as cells with either a G1S score or a G2M score greater than or equal to 0.5.

Proliferation scoring

Proliferation score of each single cell was computed using the R package UCell (*15*). Gene signature "GOBP_Cell_Division" was used by the function "AddModuleScore_UCell" to add the proliferation score for each cell. Cells with proliferation score more than the median score (0.076) were defined as proliferative cells.

Differential binding and footprint analyses

For cell type-specific differential binding and footprint analyses, snATAC-seq data from each cohort and timepoint were pooled and subset by cell types (NPCs, OPCs, and interneurons/neuroblasts). Footprint scores were calculated using the FootprintScores function from the TOBIAS toolkit (*16*). Transcription factor motifs for differential binding analysis were obtained from JASPAR (https://jaspar.elixir.no). Footprint plots were visualized using the PlotAggregate function from TOBIAS.

GEMM bulk RNA-seq sample preparation

RNA from 2000 sorted GFP+ cells from GEM models was isolated using the Absolutely RNA Nanoprep kit (Agilent, 400753-12). Immediately, cDNA synthesis was performed by using SMART-Seq v4 ultra low input RNA kit (Takara, 112219). RNA-seq libraries were prepared using the Nextera XT DNA library prep kit (Illumina, FC-131-1024). The resulting RNA-seq libraries were quantified and fragment size was determined as above and sequenced on an Illumina NextSeq 500 at the Children's Research Institute's Sequencing Facility at UT Southwestern Medical Center.

GEMM bulk RNA-seq data analysis

Sequencing adapters were trimmed using Trim Galore (v0.6.4). Trimmed reads were aligned to the mouse genome (mm10) using STAR (v2.7.3a). Mapped reads were quantified using featureCounts from the Subread package (v1.6.3). Genes with expression values of zero in more than 30% of samples were excluded from subsequent analyses. Gene expression was normalized using the voom method from the limma R package (v3.50.3), and differentially expressed genes between PIC and PC mice at 6 months were identified using the same package. Gene Set Enrichment Analysis (GSEA) of DEGs was performed using the clusterProfiler R package (v4.2.2).

TCGA RNA-seq analysis

Differential expression analysis for the TCGA GBMLGG dataset was from our previous study (*5*). TCGA RNA-seq gene expression datasets were downloaded from UCSC Xena browser (http://xena.ucsc.edu). The following TCGA studies were included: GBMLGG, CHOL, LAML, KIRP, and PCPG. $Log_2$(FPKM + 0.001) values for the interneuron lineage genes, including *GSX2*, *DLX1*, *DLX2*, *SP8*, and *SP9*, were plotted.

Cross-species transcriptomic analysis

Genes associated with the Gene Ontology (GO) biological process terms "gliogenesis" and "neurogenesis", as well as the GO molecular function term "sequence-specific DNA binding" were used to identify conserved differentially expressed genes between human IDH1/2-mutant and IDH1/2-wildtype gliomas and between cells from PIC and PC mice. Conserved differentially expressed genes with a log2(fold change) more than 1 or less than -1, and an adjusted *P* value less than 0.05 were considered significant.

<u>TCGA DNA methylation array analysis</u>
TCGA DNA methylation array datasets were downloaded from UCSC Xena browser (http://xena.ucsc.edu). The following TCGA studies were included: GBMLGG, CHOL, LAML, KIRP, and PCPG. Beta values of all probes related to the interneuron lineage genes, including *GSX2*, *DLX1*, *DLX2*, *SP8*, and *SP9* were averaged for each gene, and used to generate the plots. Beta values of all methylation probes related to *GSX2* in TCGA GBMLGG study were further extracted for the heatmap visualization.

<u>Human glioma whole genome bisulfate sequencing analysis</u>
Human glioma whole genome bisulfate sequencing (WGBS) was conducted using a previously published dataset (GSE121723) (*17*). BigWig files were downloaded from the NCBI Sequence Read Archive (SRA). The methylation levels of *GSX2* genomic region in IDH-mutant and IDH-wildtype glioma specimens, and non-malignant brain tissues were analyzed and visualized using the Integrative Genomics Viewer (IGV).

<u>Human glioma bulk ATAC-seq analysis</u>
Previously published data (GSE295378) (*18*) were used for the human glioma bulk ATAC-seq analysis. FASTQ files were downloaded from the NCBI SRA and quality-assessed using FastQC (v0.11.8). Adapters were trimmed using Trim Galore (v0.6.4) and Cutadapt (v1.9.1). Reads were aligned to the human reference genome (hg38) using Bowtie2 (v2.4.2), and the resulting SAM/BAM files were processed using SAMtools (v1.10). Genomic blacklist regions were removed from alignments using BEDtools (v2.29.2). Peak calling was performed using MACS2 (v2.2.9.1) with parameters "--nomodel --shift -100 --extsize 200" for single-end ATAC-seq data. BigWig files were generated using HOMER (v5.1) and signal tracks were visualized using IGV.

<u>Epigenome editing</u>
Mouse primary NSCs were co-transduced with an EFS-dCas9-DNMT3A3L(or dDNMT3A3L) vector (*19*) and a LentiGuide-U6-sgRNA1-U6-sgRNA2 vector to de novo methylate the *Gsx2* promoter region. *Gsx2* promoter sgRNAs were designed by using the ChopChop software (https://chopchop.cbu.uib.no). The *Hipp11* safe harbor locus targeting sgRNAs were sgRNA1 CACATAAGTTTAGACGCATT, sgRNA2 GTCATGAGACTCATACTACG. The sgRNAs used to target the *Gsx2* promoter were: sgRNA1 ACTCGGGCAGCCTATCCCGA, sgRNA2 GCTATAAAATCCTGGAGCCG.

Epigenome editing was verified by locus-specific bisulfite sequencing. Genomic DNA was extracted (DNeasy kit, Qiagen, 69506) and subjected to bisulfite conversion (EZ DNA Methylation-Lightning Kit, Zymo Research, D5030). Each conversion was split to two independent PCR reactions and the *Gsx2* promoter was then PCR amplified (NEB, M0515S) using the following primers: TGAGAGTGATGGAGGGAGAGG and TACTAAATCCCCTT CTCTCCTTCTACTA. Amplicons were gel extracted using the Gel Extraction Kit (Qiagen, 28706) and purified with the Monarch PCR & DNA Cleanup kit (NEB, T1030). Purified amplicons were sequenced by Plasmidsaurus PCR premium sequencing service. Raw sequencing reads were aligned to the amplified *Gsx2* promoter locus and the frequency of methylated to unmethylated cytosines was calculated using Bismark (V0.24.2).

For gene expression, RNA was isolated using RNeasy Plus Mini Kit (Qiagen, 74134) and subjected to cDNA synthesis using the Luna Script RT SuperMix kit (NEB, E3010). cDNA was analyzed with Luna Universal Probe qPCR Master Mix (NEB, M3004) on a Bio-Rad CFX384 real-time system with validated TaqMan qPCR probes (Thermo Fisher Scientific, *Gsx2* assay#: Mm00446650_m1; *Actb* assay#: Mm01205647_g1).

Single cell RNA-seq sample preparation

Single-cell 3' RNA-seq libraries were prepared using the 10X Genomics Chromium Next GEM Single Cell 3′ Kit v3.1 (10X Genomics, PN-1000268). Generated libraries were quantified and fragment size was determined as above. Library sequencing was performed using an Illumina NextSeq 2000 at the Children's Research Institute's Sequencing Facility at UT Southwestern Medical Center. Raw reads were generated and further processed by using 10X Genomics Cell Ranger (v8.0.1) for sample demultiplexing, barcode processing, and gene counting at the UT Southwestern BioHPC facility.

Single cell RNA-seq data analysis

Downstream analyses for single cell RNA-seq were performed using the R package Seurat (V5.3.0) (*8*). Single cells with less than 300 or more than 7,500 genes, or with greater than 12% mitochondrial fraction were filtered out for further analysis. For each sample, subsequent data processing was performed, including normalization, highly variable gene calling, scaling, cell clustering and UMAP construction. Cell type annotation was based on marker gene expression patterns as in the single nucleus multiome data analysis.

Amplicon sequencing of CRISPR Indels

To assess CRISPR-based editing of *Trp53*, *Atrx*, and *Gsx2*, nested PCR was performed using genomic DNA templates to generate amplicons centered on the PAM sites targeted by *Atrx, Trp53*, and *Gsx2* sgRNAs. The Q5 Hot Start High-Fidelity 2X Master Mix (NEB, M0494s) was used for PCR. PCR products were purified using the Monarch PCR & DNA Cleanup kit (NEB, T1030) between each PCR reaction. Final PCR products were gel-purified using the Gel Extraction Kit (Qiagen, 28706) and submitted for next-generation sequencing at the MGH DNA Core. Raw reads were analyzed using CRISPResso2 (*20*).
For *Trp53*, the following set of primers were used in sequential PCR reactions to generate amplicons for sequencing:
Outer forward *Trp53*: ATAGAGACGCTGAGTCCGGTTC
Outer reverse *Trp53*: CCTAAGCCCAAGAGGAAACAGA
Inner forward *Trp53*: TGCAGGTCACCTGTAGTGAGGTAGG
Inner reverse *Trp53*: GAAACAGGCAGAAGCTGGGGAAGAAAC
The inner PCR reaction was repeated using the above inner primers for the third PCR reaction for *Trp53*.
For *Atrx*, the following set of primers were used in sequential PCR reactions to generate amplicons for sequencing:
Outer forward *Atrx*: GCTATCTGAAACTCAATCCACG
Outer reverse *Atrx*: GACTTGGTTTCTCCTTTGCCATG

Inner forward *Atrx*: GCTTCCTGTAAGCTCATAAGTAC
Inner reverse *Atrx*: CTAATGCCATATGAGTGTAACTC
Second inner forward *Atrx*: CTCTTACATAATGGCCATTCTC
Second inner reverse *Atrx*: CTGTGAGTCATGATCATTCTTTGC
For *Gsx2*, the following set of primers were used in sequential PCR reactions to generate amplicons for sequencing:
Outer forward *Gsx2*: CGCGTTCTTTCCAAGGGTTG
Outer reverse *Gsx2*: TAACCCCTAAGTCCCTCCTCC
Inner forward *Gsx2*: AGGATGAGGACAGCGTTTACCAGCA
Inner reverse *Gsx2*: CGAAGCGCCTTTCCCCTCCTTC

*Pik3ca-H1047R* genotyping
Expression of the *Pik3ca-H1047R* allele was assessed by harvesting RNA and performing RT-PCR as previously described (*2*). RNA was isolated using the RNeasy Plus Mini Kit (Qiagen, 74134) prior to cDNA synthesis using the Luna Script RT SuperMix kit (NEB, E3010). PCR was performed using the Q5 Hot Start High-Fidelity 2X Master Mix (NEB, M0494s), and the following primers were used to detect the expression of the *Pik3ca-H1047R* transgene or *Actb*:
Forward primer *Pik3ca-H1047R*: CTAGGTAGGGGATCGGGACTCT
Reverse primer *Pik3ca-H1047R*: AATTTCTCGATTGAGGATCTTTTCT
Forward primer *Actb*: TAGGCACCAGGGTGTGATG
Reverse primer *Actb*: CATGGCTGGGGTGTTGAAGG

Immunoblotting
Cell lysates were prepared with RIPA buffer with protease inhibitor (Sigma-Aldrich, 11836153001). Protein concentrations were measured using the Protein Assay Dye Reagent (Bio-Rad, 5000006). Samples were denatured at 95 °C for 10 min, subjected to polyacrylamide gel electrophoresis (Bio-Rad), and transferred onto nitrocellulose membranes. Primary antibodies were suspended in 5% BSA in Tris-buffered saline with Tween 20 (TBST) and secondary antibodies were diluted in 5% milk in TBST. Primary antibodies used: anti-IDH1-R132H (Dianova, DIA-H09), anti-Vinculin (Sigma, V9131), anti-Cas9 (Cell Signaling, 14697S), and anti-GAPDH (Cell Signaling, 5174S). Detection was performed using Immobilon Western Chemiluminescent HRP substrate (EMD Millipore, WBKLS0500) and imaging was performed using an ImageQuant 800 biomolecular imager (Amersham).

Absolute quantification of (*R*)-2HG
Metabolites were extracted from murine glioma samples with 80% methanol in water, the supernatant was dried in a SpeedVac. The resulting pellet was resuspended in 10 μL acetonitrile with 200 ng of [$U^{13}C$]-(*R*/*S*)-2HG (Cambridge isotope laboratories, CLM-10351-PK) as internal standards, and derivatized using 90 μL of 50 mg/mL diacetyl-L-tartaric anhydride (Thermo Scientific, 336040050) in an 80:20 acetonitrile/acetic acid mixture at 75 °C for 30 min. The derivatized supernatant was dried again and reconstituted into 100 μL of 1.5 mM ammonium formate/acetonitrile (90:10) for LC/MS injection. LC/MS analysis was performed on an AB Sciex 5500 QTRAP mass spectrometer controlled by Analyst software (V1.6.1). Metabolites were separated on a Waters Acquity UPLC HSS T3 column (150 × 2.1 mm, 1.8 μM). The mobile phases were 1.5 mM ammonium formate aqueous (pH 3.6 adjusted with formic acid), and acetonitrile. Multiple reaction monitoring was used to quantify (*R*/*S*)-2HG-diacetyl tartrate

derivatives with the following transitions, R/S-2HG: m/z, 363/147, CE, −14V and [U-$^{13}$C]-R/S-2HG: m/z, 368/152, CE, −14V. Tumor volumes were calculated using tissue weights and a previously published mouse brain tissue density value (*21*). Absolute concentrations were calculated by dividing the measured (*R*)-2HG content by tumor sample volume.

Histopathology and immunohistochemistry

Histopathological analysis of brain and tumor tissues was performed by harvesting tissues and fixing immediately for 24 h in 10% formalin in PBS. Following fixation, tissues where washed and stored in 70% ethanol. Hematoxylin and eosin (H&E) staining and immunohistochemical (IHC) analyses were performed by Histowiz (histowiz.com) using a Standard Operating Procedure and fully automated workflow. Samples were processed, embedded in paraffin, and sectioned at 4 μm. Immunohistochemistry was performed on a Bond Rx autostainer (Leica Biosystems) with Heat Induced Epitope Retrieval (HIER). Antibodies used were: anti-IDH1-R132H (Dianova, DIA-H09), anti-SOX2 (Cell Signaling, 14962), and anti-GFP (Abcam, ab183734). Chromagen development was done using the Bond Polymer Refine Detection Kit (Leica Biosystems) which was used according to the manufacturer's protocol. After staining, sections were dehydrated and film coverslipped using a TissueTek-Prisma Coverslipper. Whole slide scanning (40X) was performed on an Aperio AT2 (Leica Biosystems).

Magnetic resonance imaging

Magnetic resonance imaging (MRI) was performed at the UT Southwestern Preclinical PET/MRI core using a 9.4T Bruker BioSpec scanner equipped with an MRI CryoProbe, a cryogenically cooled, anatomically shaped 2×2 receive-only coil. Mice were anesthetized with 1.5% isoflurane, and their heads were fixed using tooth and ear bars. A close warm-water circuit maintained the body temperature at 37 °C. T2-weighted, multi-slice spin echo (RARE: Rapid Acquisition with Relaxation Enhancement) imaging was performed. Imaging parameters included: field of view = 20x15 mm$^2$, in-plane resolution = 78x78 μm$^2$ slice thickness = 0.5 mm, matrix = 256x192, RARE factor = 8, TEeff/TR = 35/2500 ms, and the number of averages = 5. The total scan time was 5 minutes.

Mouse methylation array analysis

Tumor bearing mice (confirmed by MRI) and normal healthy mice were euthanized, cardiac perfusion was performed and whole brain was extracted. GFP+ glioma tissues were visualized using a fluorescence flashlight system (Nightsea, Xite-RB-GO) and carefully dissected. GFP+ tumor samples and normal brain tissues were snap frozen in liquid nitrogen and kept in -80 °C until further processing. Methylation array analysis was conducted at the Genomics & Microarray core at UT Southwestern Medical Center using a standardized protocol. Genomic DNA was extracted (DNeasy kit, Qiagen, 69506) and subjected to bisulfite conversion (EZ-96 DNA Methylation Kit, Zymo Research, D5004). The DNA methylation status of the bisulfite converted DNA was obtained using the Infinium Mouse Methylation BeadChip (Illumina). Raw signal intensities were assessed and analyzed with GenomeStudio Methylation Module (Illumina). Methylation probes with mean beta values more than 0.1 in normal brain tissues from C mice (harboring no mutations, expressing only Cas9-2A-GFP) and standard deviation greater than 0.1 in glioma samples from PIC and PC mice were used to construct DNA methylation histograms.

Whole brain GFP imaging

Mice were euthanized, cardiac perfusion with PBS was performed, and whole brains were extracted. Immediately, mouse whole brains were fixed at room temperature in 10% formalin in PBS for 24 hours, then stored in PBS with 0.02% $NaN_3$ at 4 °C until embedding. Whole brains were embedded in agarose (Sigma, A0169) following established procedures (*22*). After overnight crosslinking in sodium borohydride, the agarose blocks were transferred to phosphate buffer (PB) for storage at 4°C until TissueCyte imaging. Whole brains were subjected to serial two-photon tomography (STPT) on the TissueCyte 1000 (TissueVision) using established procedures (*22*) in the UT Southwestern Whole Brain Microscopy Facility (RRID:SCR_017949). The acquired raw image tiles were first trimmed and subjected to flat field correction, then stitched into 2D mosaic coronal section images using the Autostitcher software (TissueVision).

Statistical analysis

Statistical analysis was performed using Graphpad Prism software. Unless otherwise specified in the figure legend, *P* values were calculated by unpaired *t*-tests for tests of statistical significance involving comparison of two groups or by one-way ANOVA nonparametric tests for three groups. For survival study, *P* value was calculated by log-rank test. For cell states comparison between groups, *P* value was calculated by Chi-square test.

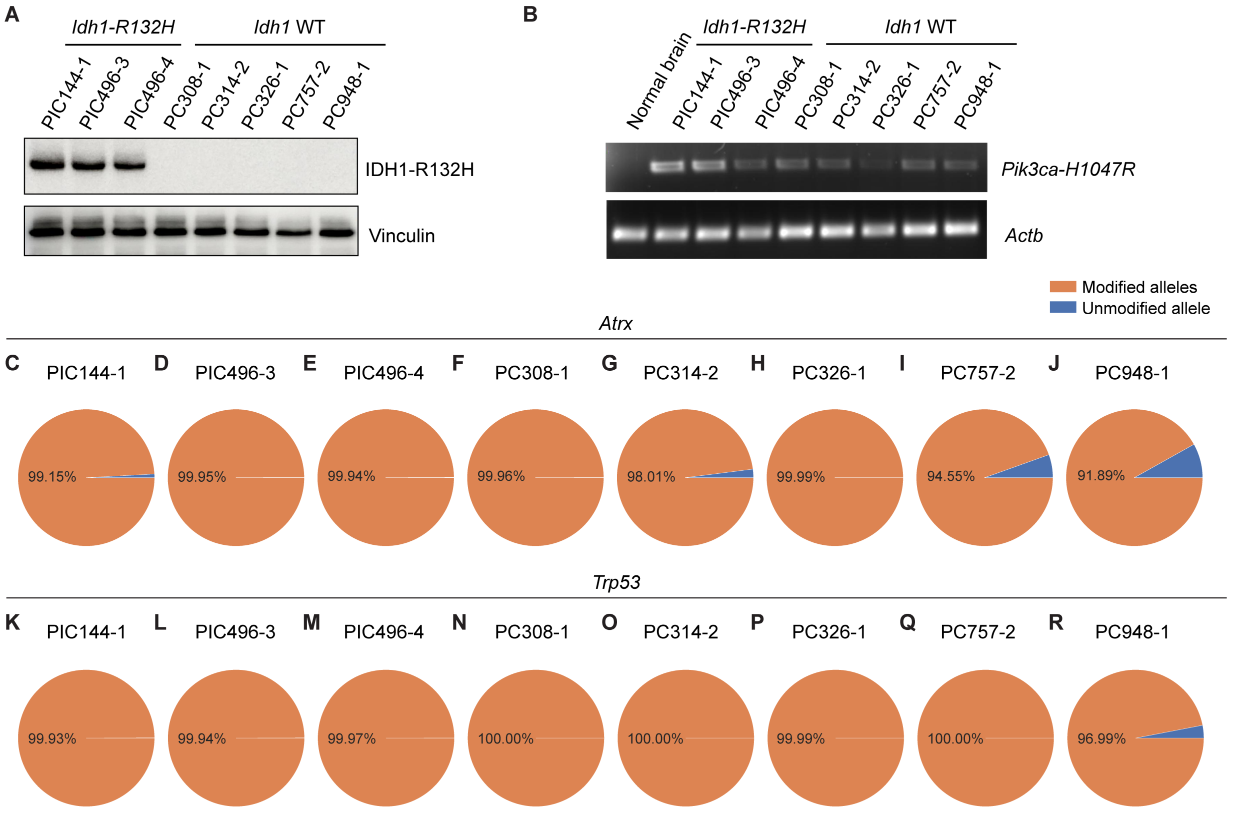


**Fig. S1. Validation of genetic alterations in engineered astrocytomas.**

(**A**) Immunoblot of IDH1-R132H oncoprotein expression, (**B**) RT-PCR assays for *Pik3ca-H1047R* or *Actb* expression, and (**C**)-(**R**) Amplicon sequencing of genomic DNA regions of *Atrx* (C)-(I) and *Trp53* (K)-(R) targeted by CRISPR sgRNAs in PIC and PC glioma-derived neurosphere lines. In (B), brain tissue from a naïve mouse was used as a negative control. In all panels, numbers following genotype abbreviations (e.g. PICXXX-X) represent unique identifiers for individual mice.

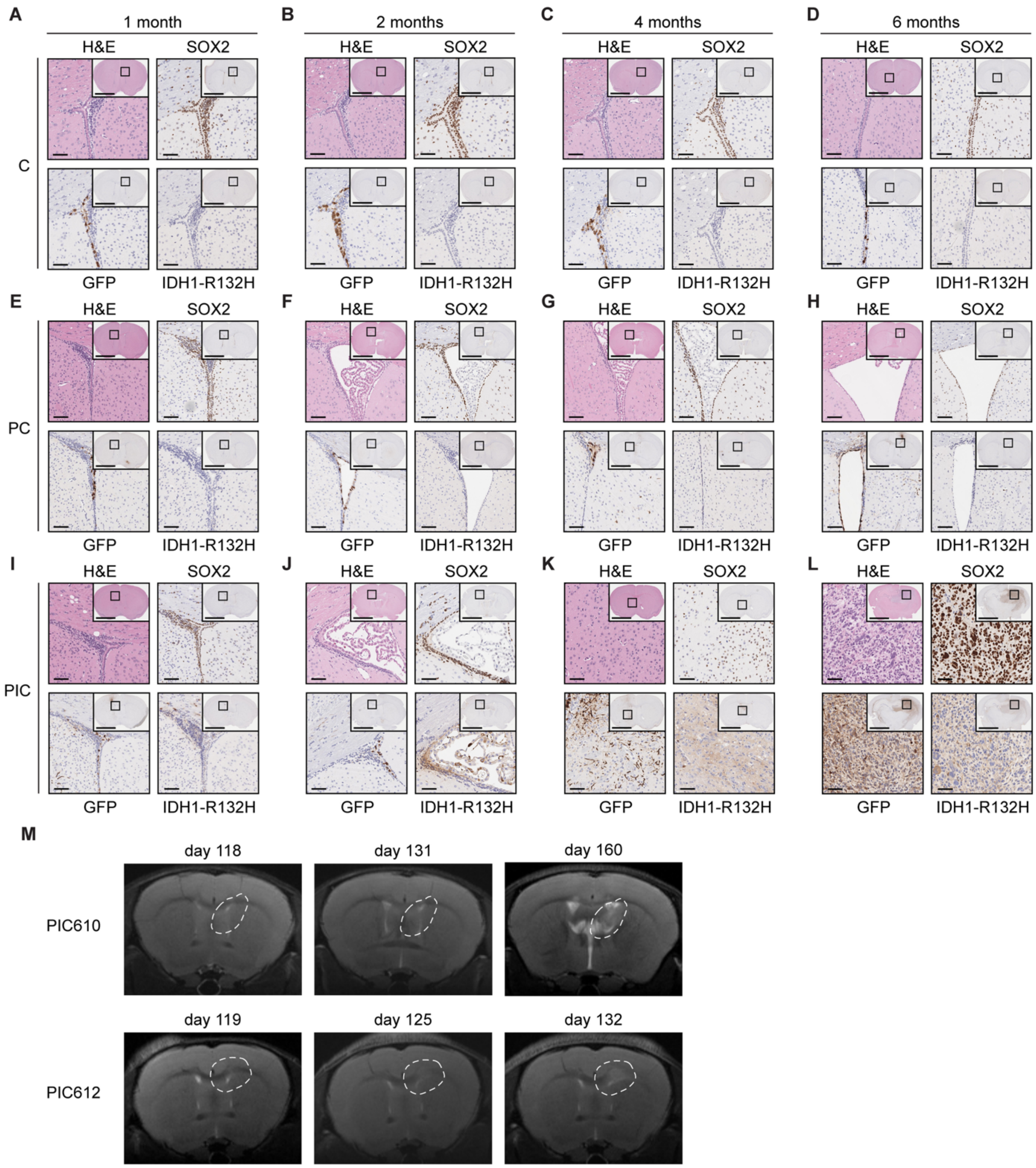


**Fig. S2. Longitudinal histopathological and imaging analyses of genetically engineered mice.**

(**A**)-(**L**) H&E, SOX2, GFP, and IDH1-R132H IHC analyses of representative C (A)-(D), PC (E)-(H), and PIC (I)-(L) GEM mice at 1, 2, 4, and 6 months after AAV injection. (**M**) Brain MRI images show spatial and temporal dynamics of glioma initiation in PIC mice. Scale bars in main images and insets in (A)-(L) indicate 100 μm and 3 mm, respectively. In (M), days reflect ages of mice and dashed lines highlight regions of tumor development (evident at last time point).

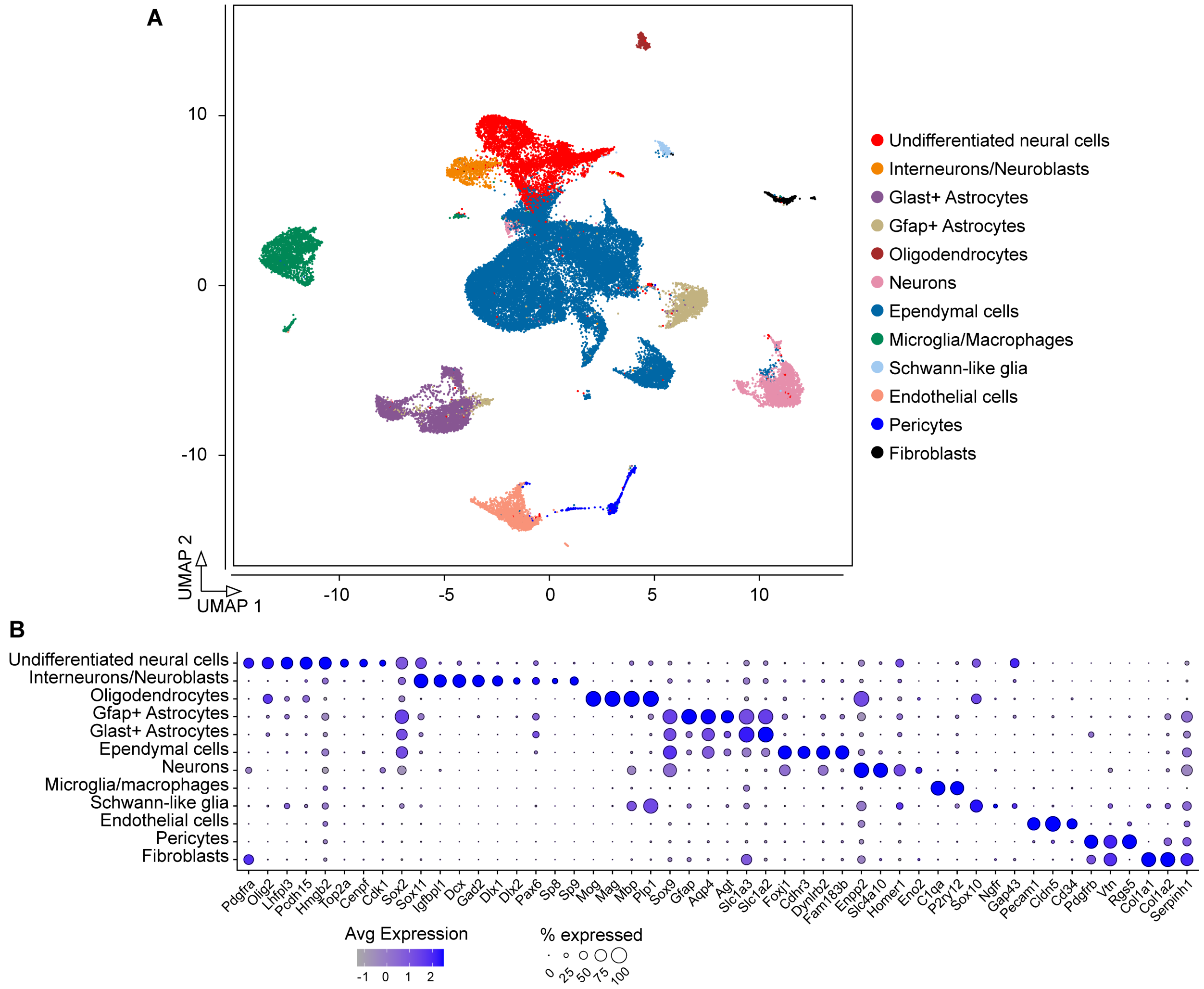


**Fig. S3. Single cell genomics analyses identify distinct neural cell populations in genetically engineered mouse brains.**

(**A**) UMAP plot showing identified neural cell populations in C, PC, and PIC mice using single cell genomics. (**B**) Dotplot showing the expression patterns of select cell type-specific markers across brain-resident cell populations.

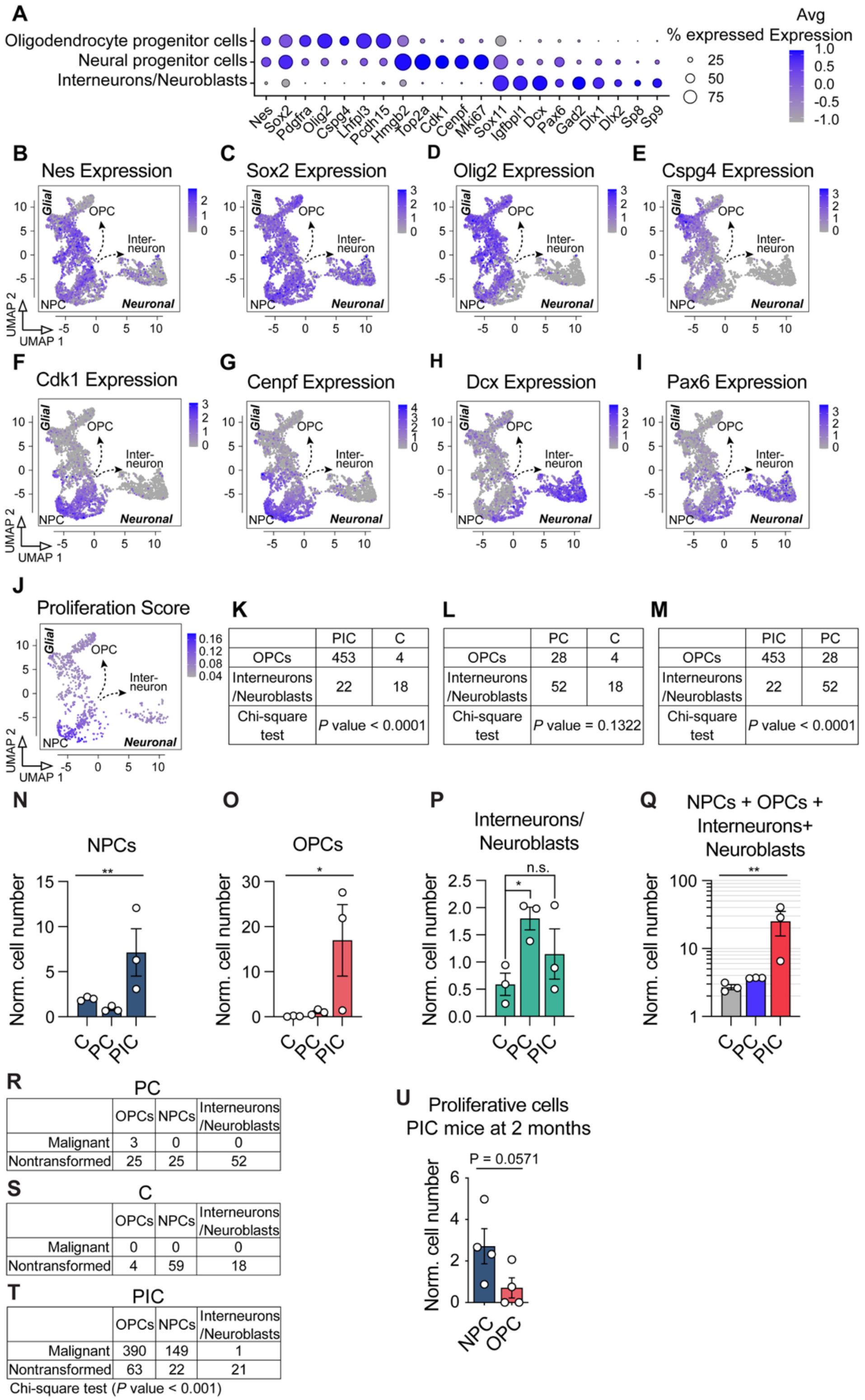


**Fig. S4. Mutant IDH activates NPCs and alters their lineage specification.**

(**A**) Dotplot showing the expression pattern of selected cell type specific canonical markers for oligodendrocyte progenitor cells (OPCs), neural progenitor cells (NPCs), and interneurons/neuroblasts. (**B**)-(**I**) UMAP overlays of expression of selected canonical marker genes. (**J**) UMAP overlays of proliferation scores to mark proliferating cells in OPCs, NPCs and interneurons/neuroblasts from C, PC, and PIC mice at 4 months. (**K**)-(**M**) Comparisons of absolute numbers of OPCs and interneurons/neuroblasts in PIC vs C (K), PC vs C (L), and PIC vs PC (M) at 4 months. (**N**)-(**Q**) Quantification of NPCs (N), OPCs (O), interneurons/neuroblasts (P), and all states combined (Q) in C, PC, and PIC mice at 4 months. Cell numbers were normalized (Norm.) by calculating their incidence per 100 cells sequenced in $GFP^+$ populations. (**R**)-(**T**) Comparisons of absolute numbers of malignant and nontransformed cells in OPCs, NPCs and interneurons/neuroblasts in PC (R), C (S), and PIC (T) mice at 4 months. (**U**) Quantification of proliferative NPCs and OPCs in PIC mice at 2 months. In (K)-(U), cells are derived from ≥3 mice at each time point in each cohort. In (N)-(Q), data are means ± SEM; *P* values in (N), (O), and (Q) were calculated by one-way ANOVA nonparametric tests, *P* values in (P) were calculated by unpaired t-tests. * $P < 0.05$, ** $P < 0.01$, *** $P < 0.001$, n.s. = not significant. In (U), *P* value was calculated by an unpaired t-test.

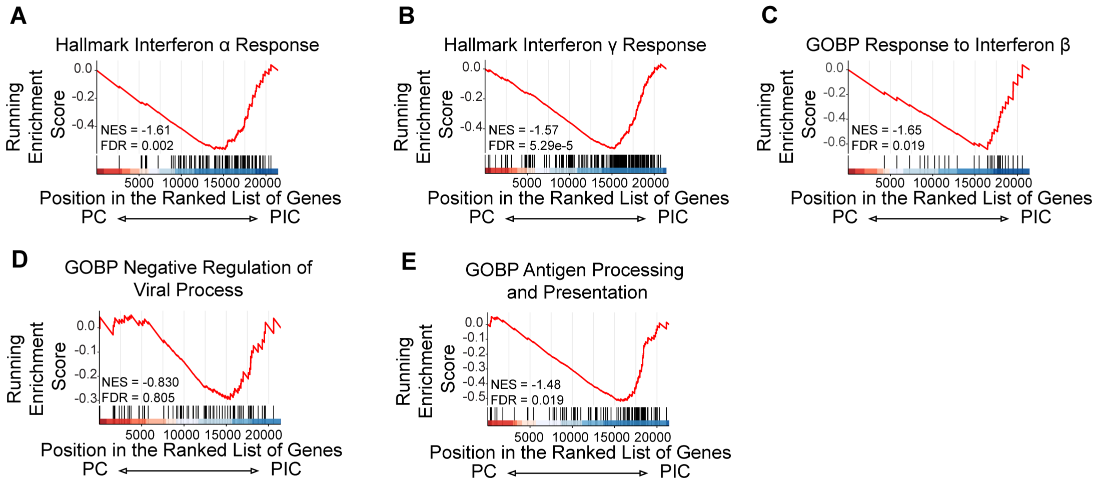


**Fig. S5. Mutant IDH1 suppresses interferon signaling in PIC mice.**

Selected GSEA plots depicting expression of interferon-related gene sets in engineered cells from PIC and PC mice. (**A**) Hallmark interferon α response. (**B**) Hallmark interferon γ response. (**C**) Gene Ontology (GO) biological process (BP) response to interferon β. (**D**) GOBP negative regulation of viral process. (**E**) GOBP antigen processing and presentation.

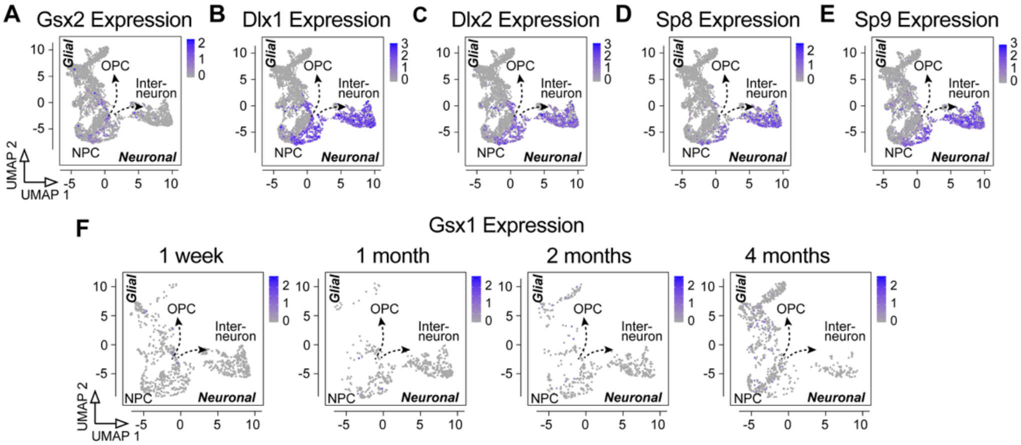


**Fig. S6. Expression of interneuron developmental genes in select neural cell populations.** (**A**)-(**E**) UMAP overlays of expression of neuroblast/interneuron lineage genes in NPCs, OPCs and interneurons/neuroblasts from C, PC, and PIC mice. (**F**) UMAP overlays of *Gsx1* expression in NPCs, OPCs and interneurons/neuroblasts from C, PC, and PIC mice during gliomagenesis.

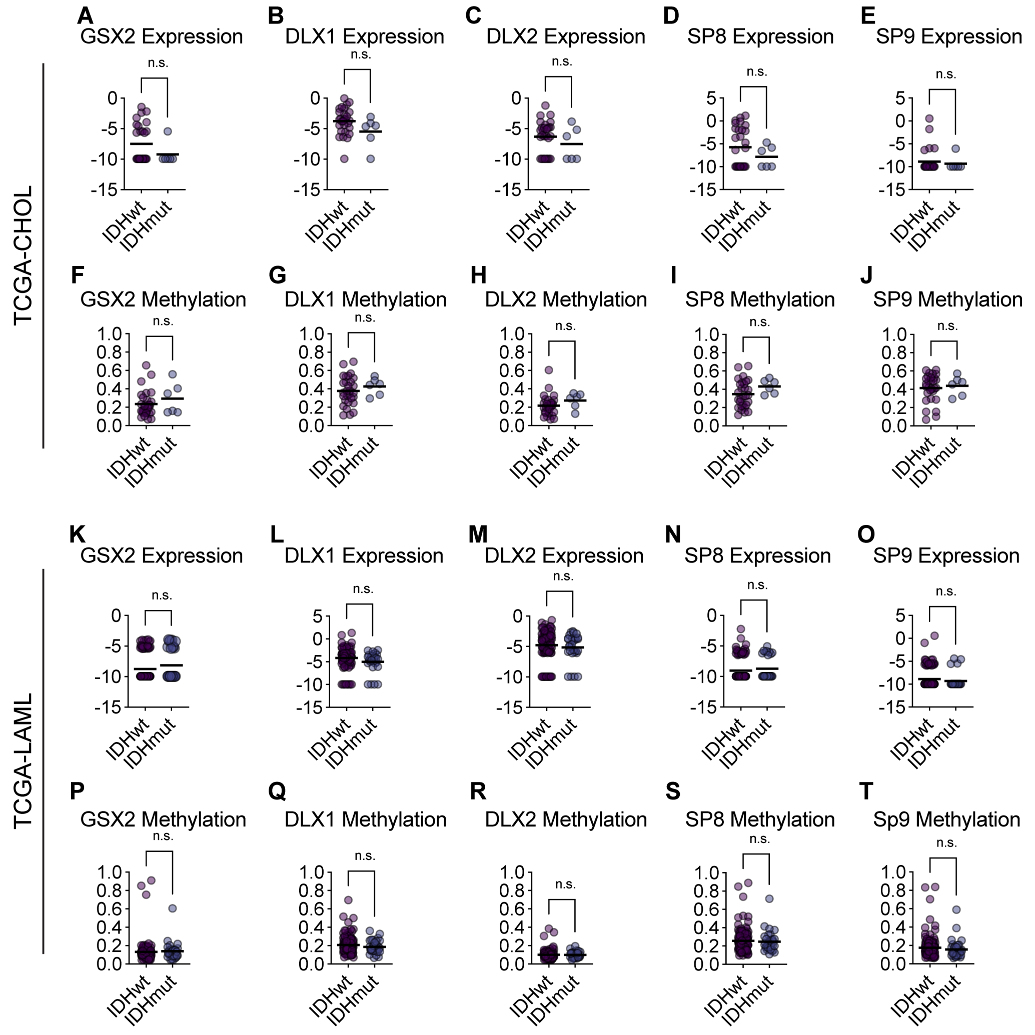


**Fig. S7. Expression and methylation of interneuron developmental genes in other IDH-mutant cancers.**

(**A**)-(**J**) Neuroblast/interneuron lineage gene expression (A)-(E) and methylation (F)-(J) in IDHmut vs IDHwt human cholangiocarcinoma samples. (**K**)-(**T**) Neuroblast/interneuron lineage gene expression (K)-(O) and methylation (P)-(T) in IDHmut vs IDHwt human acute myeloid leukemia samples. In (A)-(E) and (K)-(O), values are $\log_2$(FPKM + 0.001). In (F)-(J) and (P)-(T), values are beta values. In all panels, bars are median values; *P* values were calculated by unpaired t-tests, n.s. = not significant.

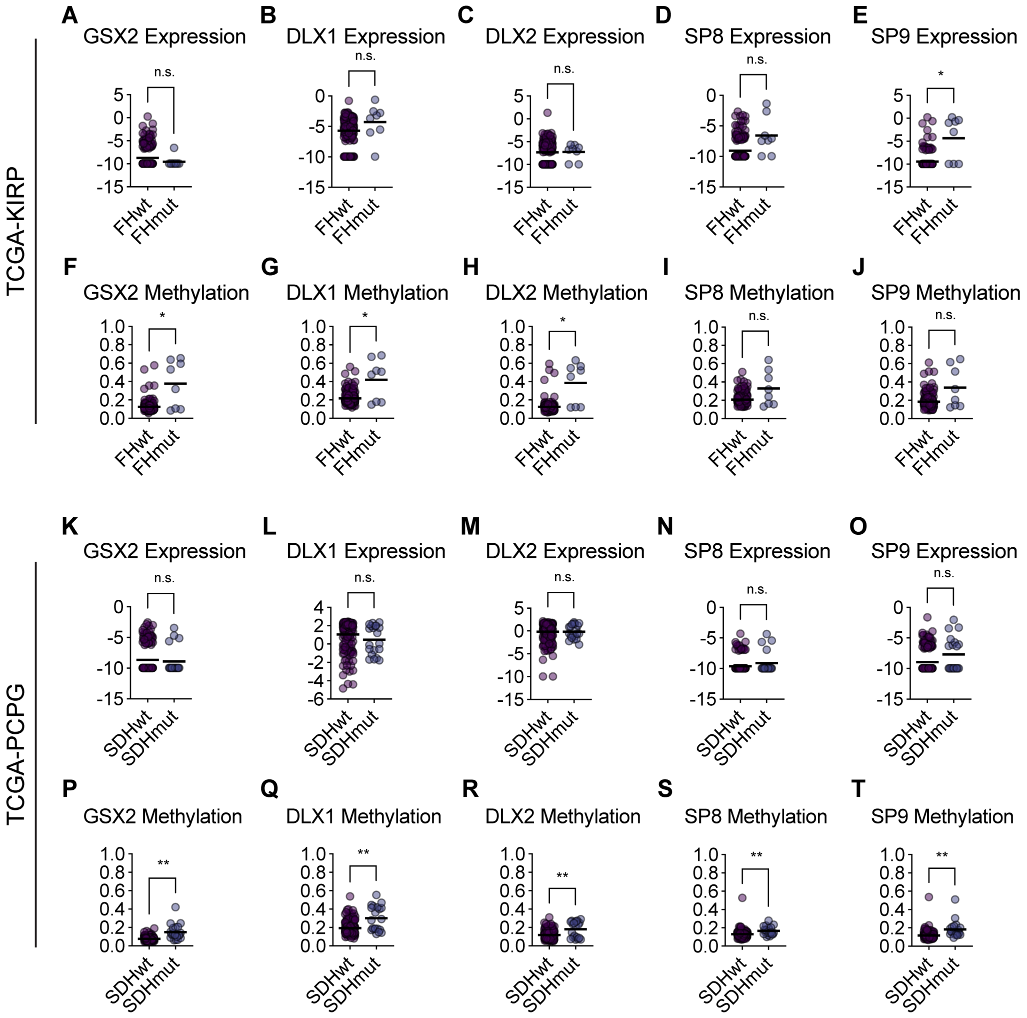


**Fig. S8. Expression and methylation of interneuron developmental genes in SDH- and FH-mutant cancers.**

(**A**)-(**J**) Neuroblast/interneuron lineage gene expression (A-E) and methylation (F-J) in FHmut vs FHwt human papillary renal cell carcinoma samples. (**K**)-(**T**) Neuroblast/interneuron lineage gene expression (K-O) and methylation (P-T) in SDHmut vs SDHwt human pheochromocytoma and paraganglioma samples. In (A)-(E) and (K)-(O), values are $\log_2$(FPKM + 0.001). In (F)-(J) and (P)-(T), values are beta values. In all panels, bars are median values, $P$ values were calculated by unpaired t-tests, * $P < 0.05$, ** $P < 0.01$, n.s. = not significant.

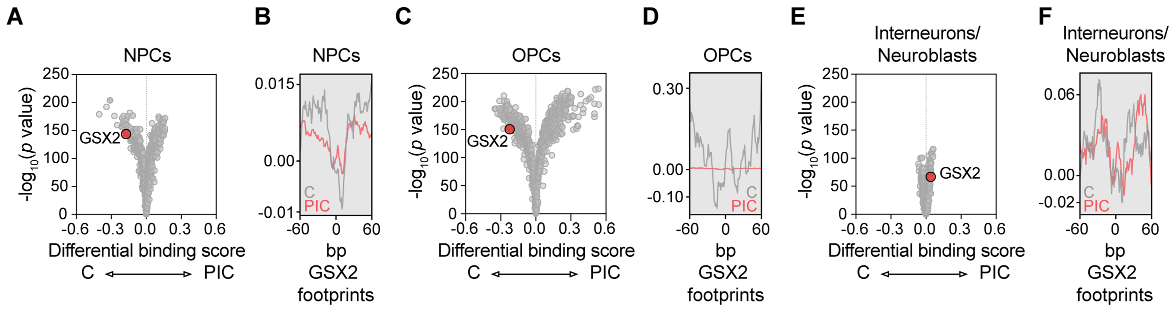


**Fig. S9. Mutant IDH inhibits GSX2 chromatin occupancy in NPCs and OPCs.**

(**A**)-(**F**) Comparison of genome-wide DNA binding scores for transcription factors in NPCs (A)-(B), OPCs (C)-(D), and interneurons/neuroblasts (E)-(F) from PIC and C mice at 4 months. In (A), (C), and (E), volcano plots are shown, with the TOBIAS differential binding score on the x-axis and $-\log_{10}$(p value) on the y-axis. In (B), (D), and (F), footprint plot overlays are shown for *Gsx2* in cells from PIC (red line) and C (grey line) mice.

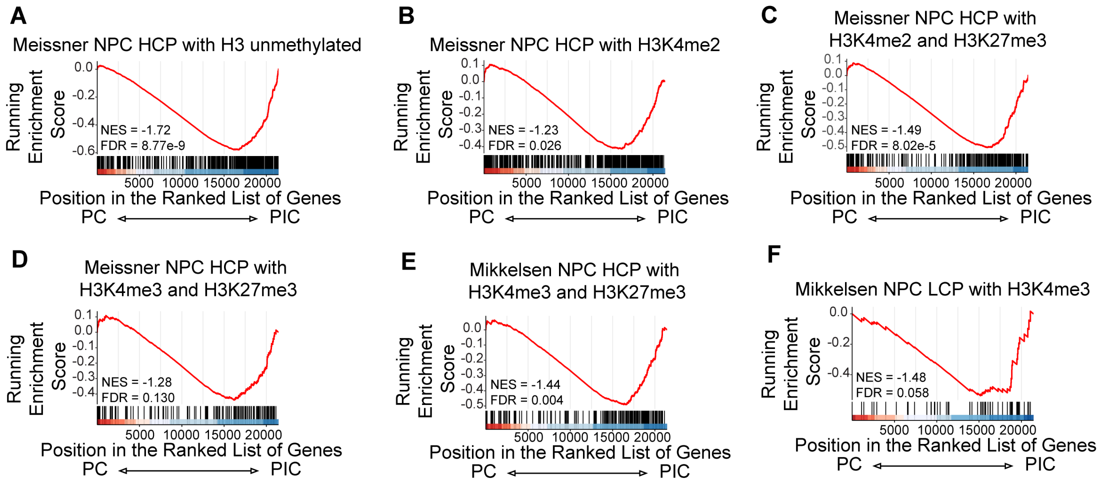


**Fig. S10. Mutant IDH suppresses genes with high CpG density.**

Selected GSEA plots depicting expression of high CpG content (HCP) and low CpG content (LCP) gene sets in engineered cells from PIC and PC mice. (**A**) Meissner NPC HCP with H3 unmethylated. (**B**) Meissner NPC HCP with H3K4me2. (**C**) Meissner NPC HCP with H3K4me2 and H3K27me3. (**D**) Meissner NPC HCP with H3K4me3 and H3K27me3. (**E**) Mikkelsen NPC HCP with H3K4me3 and H3K27me3. (**F**) Mikkelsen NPC LCP with H3K4me3.

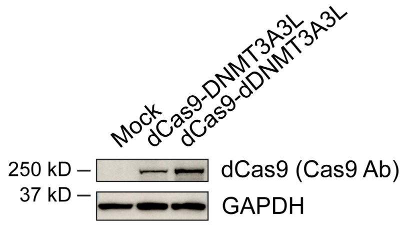


**Fig. S11. Validation of dCas9-(d)DNMT3A3L fusion protein expression constructs.** Immunoblot validation of dCas9-(d)DNMT3A3L fusion protein expression in HEK293 cells using an anti-Cas9 antibody. Fusion proteins appear as bands around 225 kDa, larger than the expected ~158 kDa for unmodified dCas9. GAPDH serves as a loading control.

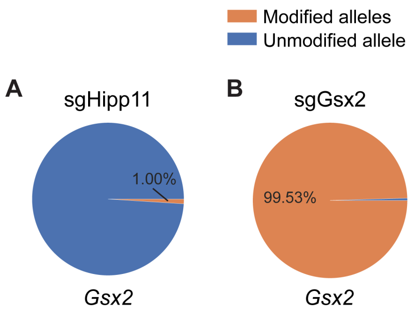


**Fig. S12. Amplicon sequencing of CRISPR indels in the *Gsx2* gene.**

(**A**) and (**B**) Amplicon sequencing of the genomic DNA region targeted by the *Gsx2* CRISPR sgRNA in GFP$^+$ neural cells. Cells were recovered from PC mice injected with AAV particles encoding an sgRNA targeting the (A) *Hipp11* safe harbor locus or (B) *Gsx2*.

### References

1. R. J. Platt, S. Chen, Y. Zhou, M. J. Yim, L. Swiech, H. R. Kempton, J. E. Dahlman, O. Parnas, T. M. Eisenhaure, M. Jovanovic, D. B. Graham, S. Jhunjhunwala, M. Heidenreich, R. J. Xavier, R. Langer, D. G. Anderson, N. Hacohen, A. Regev, G. Feng, P. A. Sharp, F. Zhang, CRISPR-Cas9 knockin mice for genome editing and cancer modeling. *Cell* **159**, 440–455 (2014).

2. J. R. Adams, K. Xu, J. C. Liu, N. M. Agamez, A. J. Loch, R. G. Wong, W. Wang, K. L. Wright, T. F. Lane, E. Zacksenhaus, S. E. Egan, Cooperation between Pik3ca and p53 mutations in mouse mammary tumor formation. *Cancer Res.* **71**, 2706–2717 (2011).

3. M. Sasaki, C. B. Knobbe, J. C. Munger, E. F. Lind, D. Brenner, A. Brüstle, I. S. Harris, R. Holmes, A. Wakeham, J. Haight, A. You-Ten, W. Y. Li, S. Schalm, S. M. Su, C. Virtanen, G. Reifenberger, P. S. Ohashi, D. L. Barber, M. E. Figueroa, A. Melnick, J.-C. Zúñiga-Pflücker, T. W. Mak, IDH1(R132H) mutation increases murine haematopoietic progenitors and alters epigenetics. *Nature* **488**, 656–659 (2012).

4. M. Sasaki, C. B. Knobbe, M. Itsumi, A. J. Elia, I. S. Harris, I. I. C. Chio, R. A. Cairns, S. McCracken, A. Wakeham, J. Haight, A. Y. Ten, B. Snow, T. Ueda, S. Inoue, K. Yamamoto, M. Ko, A. Rao, K. E. Yen, S. M. Su, T. W. Mak, D-2-hydroxyglutarate produced by mutant IDH1 perturbs collagen maturation and basement membrane function. *Genes Dev* **26**, 2038–2049 (2012).

5. D. D. Shi, M. R. Savani, M. M. Levitt, A. C. Wang, J. E. Endress, C. E. Bird, J. Buehler, S. A. Stopka, M. S. Regan, Y.-F. Lin, V. T. Puliyappadamba, W. Gao, J. Khanal, L. Evans, J. H. Lee, L. Guo, Y. Xiao, M. Xu, B. Huang, R. B. Jennings, D. M. Bonal, M. S. Martin-Sandoval, T. Dang, L. C. Gattie, A. B. Cameron, S. Lee, J. M. Asara, H. I. Kornblum, T. W. Mak, R. E. Looper, Q.-D. Nguyen, S. Signoretti, S. Gradl, A. Sutter, M. Jeffers, A. Janzer, M. A. Lehrman, L. G. Zacharias, T. P. Mathews, J.-A. Losman, T. E. Richardson, D. P. Cahill, R. J. DeBerardinis, K. L. Ligon, L. Xu, P. Ly, N. Y. R. Agar, K. G. Abdullah, I. S. Harris, W. G. Kaelin Jr., S. K. McBrayer, De novo pyrimidine synthesis is a targetable vulnerability in IDH mutant glioma. *Cancer Cell* **40**, 939-956.e16 (2022).

6. C. Lothian, N. Prakash, U. Lendahl, G. M. Wahlström, Identification of both general and region-specific embryonic CNS enhancer elements in the nestin promoter. *Exp Cell Res* **248**, 509–519 (1999).

7. R. C. Challis, S. Ravindra Kumar, K. Y. Chan, C. Challis, K. Beadle, M. J. Jang, H. M. Kim, P. S. Rajendran, J. D. Tompkins, K. Shivkumar, B. E. Deverman, V. Gradinaru, Systemic AAV vectors for widespread and targeted gene delivery in rodents. *Nat. Protoc.* **14**, 379–414 (2019).

8. Y. Hao, T. Stuart, M. H. Kowalski, S. Choudhary, P. Hoffman, A. Hartman, A. Srivastava, G. Molla, S. Madad, C. Fernandez-Granda, R. Satija, Dictionary learning for integrative, multimodal and scalable single-cell analysis. *Nature Biotechnology* **42**, 293–304 (2024).

9. T. Stuart, A. Srivastava, S. Madad, C. A. Lareau, R. Satija, Single-cell chromatin state analysis with Signac. *Nature Methods* **18**, 1333–1341 (2021).

10. R. Gao, S. Bai, Y. C. Henderson, Y. Lin, A. Schalck, Y. Yan, T. Kumar, M. Hu, E. Sei, A. Davis, F. Wang, S. F. Shaitelman, J. R. Wang, K. Chen, S. Moulder, S. Y. Lai, N. E. Navin, Delineating copy number and clonal substructure in human tumors from single-cell transcriptomes. *Nature Biotechnology* **39**, 599–608 (2021).

11. C. Trapnell, D. Cacchiarelli, J. Grimsby, P. Pokharel, S. Li, M. Morse, N. J. Lennon, K. J. Livak, T. S. Mikkelsen, J. L. Rinn, The dynamics and regulators of cell fate decisions are revealed by pseudotemporal ordering of single cells. *Nature Biotechnology* **32**, 381–386 (2014).

12. I. Tirosh, A. S. Venteicher, C. Hebert, L. E. Escalante, A. P. Patel, K. Yizhak, J. M. Fisher, C. Rodman, C. Mount, M. G. Filbin, C. Neftel, N. Desai, J. Nyman, B. Izar, C. C. Luo, J. M. Francis, A. A. Patel, M. L. Onozato, N. Riggi, K. J. Livak, D. Gennert, R. Satija, B. V. Nahed, W. T. Curry, R. L. Martuza, R. Mylvaganam, A. J. Iafrate, M. P. Frosch, T. R. Golub, M. N. Rivera, G. Getz, O. Rozenblatt-Rosen, D. P. Cahill, M. Monje, B. E. Bernstein, D. N. Louis, A. Regev, M. L. Suva, Single-cell RNA-seq supports a developmental hierarchy in human oligodendroglioma. *Nature* **539**, 309–313 (2016).

13. A. Spitzer, S. Gritsch, M. Nomura, A. Jucht, J. Fortin, R. Raviram, H. R. Weisman, L. N. Gonzalez Castro, N. Druck, R. Chanoch-Myers, J. J. Y. Lee, R. Mylvaganam, R. Lee Servis, J. M. Fung, C. K. Lee, H. Nagashima, J. J. Miller, I. Arrillaga-Romany, D. N. Louis, H. Wakimoto, W. Pisano, P. Y. Wen, T. W. Mak, M. Sanson, M. Touat, D. A. Landau, K. L. Ligon, D. P. Cahill, M. L. Suvà, I. Tirosh, Mutant IDH inhibitors induce lineage differentiation in IDH-mutant oligodendroglioma. *Cancer Cell* **42**, 904-914.e9 (2024).

14. A. S. Venteicher, I. Tirosh, C. Hebert, K. Yizhak, C. Neftel, M. G. Filbin, V. Hovestadt, L. E. Escalante, M. L. Shaw, C. Rodman, S. M. Gillespie, D. Dionne, C. C. Luo, H. Ravichandran, R. Mylvaganam, C. Mount, M. L. Onozato, B. V. Nahed, H. Wakimoto, W. T. Curry, A. J. Iafrate, M. N. Rivera, M. P. Frosch, T. R. Golub, P. K. Brastianos, G. Getz, A. P. Patel, M. Monje, D. P. Cahill, O. Rozenblatt-Rosen, D. N. Louis, B. E. Bernstein, A. Regev, M. L. Suva, Decoupling genetics, lineages, and microenvironment in IDH-mutant gliomas by single-cell RNA-seq. *Science* **355**, eaai8478 (2017).

15. M. Andreatta, S. J. Carmona, UCell: Robust and scalable single-cell gene signature scoring. *Computational and Structural Biotechnology Journal* **19**, 3796–3798 (2021).

16. M. Bentsen, P. Goymann, H. Schultheis, K. Klee, A. Petrova, R. Wiegandt, A. Fust, J. Preussner, C. Kuenne, T. Braun, J. Kim, M. Looso, ATAC-seq footprinting unravels kinetics of transcription factor binding during zygotic genome activation. *Nature Communications* **11**, 4267 (2020).

17. Y. Wu, M. Fletcher, Z. Gu, Q. Wang, B. Costa, A. Bertoni, K. H. Man, M. Schlotter, J. Felsberg, J. Mangei, M. Barbus, A. C. Gaupel, W. Wang, T. Weiss, R. Eils, M. Weller, H. Liu, G. Reifenberger, A. Korshunov, P. Angel, P. Lichter, C. Herrmann, B. Radlwimmer,

Glioblastoma epigenome profiling identifies SOX10 as a master regulator of molecular tumour subtype. *Nat. Commun.* **11**, 6434 (2020).

18. M. C. Garrett, R. Albano, T. Carnwath, S. Shah, D. Woo, M. Lamba, D. R. Plas, A. Paranjpe, K. Roskin, C. Zhao, R. Lu, Chromatin structure predicts survival in glioma patients. *Scientific Reports* **12**, 8221 (2022).

19. R. S. Guerra-Resendez, S. L. Lydon, A. J. Ma, G. C. Bedford, D. R. Reed, S. Kim, E. R. Teran, T. Nishiguchi, M. Escobar, A. R. DiNardo, I. B. Hilton, Characterization of Rationally Designed CRISPR/Cas9-Based DNA Methyltransferases with Distinct Methyltransferase and Gene Silencing Activities in Human Cell Lines and Primary Human T Cells. *ACS Synth. Biol.* **14**, 384–397 (2025).

20. K. Clement, H. Rees, M. C. Canver, J. M. Gehrke, R. Farouni, J. Y. Hsu, M. A. Cole, D. R. Liu, J. K. Joung, D. E. Bauer, L. Pinello, CRISPResso2 provides accurate and rapid genome editing sequence analysis. *Nat Biotechnol* **37**, 224–226 (2019).

21. H. W. Bothe, W. Bodsch, K. A. Hossmann, Relationship between specific gravity, water content, and serum protein extravasation in various types of vasogenic brain edema. *Acta Neuropathol* **64**, 37–42 (1984).

22. D. M. O. Ramirez, A. D. Ajay, M. P. Goldberg, J. P. Meeks, "Serial Multiphoton Tomography and Analysis of Volumetric Images of the Mouse Brain" in *Multiphoton Microscopy*, E. Hartveit, Ed. (Springer New York, New York, NY, 2019; https://doi.org/10.1007/978-1-4939-9702-2_9), pp. 195–224.

**Data S1. DNA methylation array analysis of brain tissues from PIC, PC, and C mice. (separate file)**

This file contains the DNA methylation array analysis using primary tumor tissues from PIC and PC mice and normal brain samples from C mice.

**Data S2. RNA-seq results comparing PIC and PC mouse tissues. (separate file)**

This file contains the results of RNA-seq and differential gene expression analyses of PIC and PC mouse tissues at 6 months. PC mice without overt tumors at 6 months were chosen for sequencing to facilitate identification of gene expression patterns associated with mutant Idh-dependent mechanisms of transformation.

**Data S3. Gene set enrichment analysis of engineered cells from PIC and PC mice. (separate file)**

This file contains the GSEA results of all pathways using gene expression data from engineered cells recovered from PIC and PC mice.

**Data S4. Cross-species gene expression analysis in IDH-mutant and IDH-wildtype brain specimens. (separate file)**

This file contains the results of conserved differentially expressed genes related to gliogenesis, neurogenesis or transcription factors in human and mouse IDH-mutant and IDH-wildtype glioma samples.

**Data S5. Differential transcription factor binding analysis of engineered cells from PIC, PC and C mice. (separate file)**

This file contains the differential binding analysis of all transcription factors using snATAC-seq data of NPCs, OPCs, and interneurons/neuroblasts at 4 months from PIC, PC, and C mice. Analyses compare cells from PIC versus PC mice, as well as cells from PIC versus C mice.
